# Blood pressure pulsations modulate olfactory bulb neuronal activity via mechanosensitive ion channels

**DOI:** 10.1101/2023.07.27.550787

**Authors:** Luna Jammal Salameh, Sebastian H. Bitzenhofer, Ileana L. Hanganu-Opatz, Mathias Dutschmann, Veronica Egger

## Abstract

The transmission of heartbeat through the cerebral vascular system is known to cause intracranial pressure pulsations. Here we describe that arterial pressure pulsations within the brain can directly modulate central neuronal activity. In a semi-intact rat brain preparation, pressure pulsations elicit correlated local field oscillations in the olfactory bulb (OB) that are sensitive to hypoxia and block of mechanosensitive channels. We find that mitral cell spiking activity is in part synchronized to these oscillations. Indeed, in awake animals the firing of a subset of OB neurons is entrained to heartbeat within ∼ 20 ms. Several lines of evidence indicate the expression of a mechanosensitive ion channel within the mitral cell membranes, most likely Piezo2, implementing a pressure pulsation transduction pathway and thus baroreception within the OB. We propose that this intrinsic interoceptive mechanism modulates OB neuronal activity e.g. during arousal and also could influence brain activity on a wider scale.

## 1. Introduction

Both spontaneous and sensory-evoked neuronal network oscillations are a hallmark of olfactory systems across phylae (Kay, 2015). In order to investigate the origins and mechanisms of these oscillations within the olfactory bulb, we had developed a semi-intact preparation of the rat olfactory system (nose brain preparation, NBP, Pérez de los Cobos Pallarés et al., 2015). In this type of preparations, the tissue of interest – as in other *in situ* preparations of the brainstem (Paton et al., 2022) or the heart (Bell et al., 2011) – is perfused using a peristaltic pump (Fig. 1A). In our system, this pump generates pressure pulsations within the vascular system that with regards to both the amplitude and time course are likely to fit within the physiological range of heart-beat induced pulsations in the intracranial pressure *in vivo* (see Wagshul et al., 2011).

**Figure 1:**
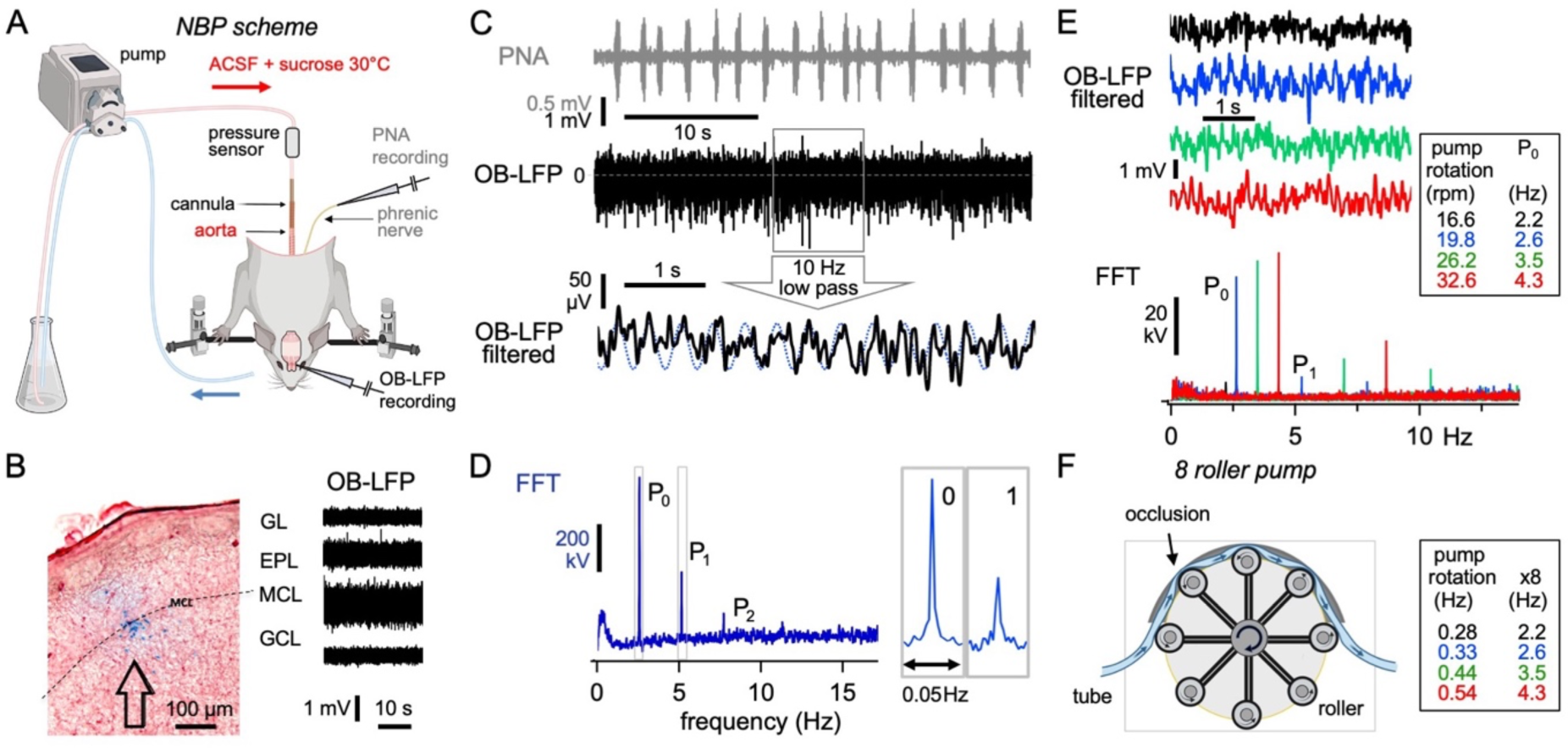
Slow oscillations in the olfactory bulb LFP in a semi-intact perfused brain preparation due to peristaltic pump roller movements **A** Schematic depiction of the nose brain preparation. The decerebrated preparation is perfused via the cannulated descending aorta. Spontaneous bulbar LFPs and phrenic nerve activity (PNA) are recorded in parallel. Created with www.Biorender.com. **B** Recording location of the LFP within the OB. Left panel: Staining of recording electrode location (see Methods). Right panel: LFP activity is highest in the mitral cell layer. **C** Exemplary recording of rhythmic PNA and OB-LFP within the mitral cell layer. Bottom trace: low-pass filtering of the LFP reveals a slow oscillatory component. **D** Lower end of the Fourier spectrum of the above OB-LFP recording. Note the harmonic peaks P_1_, P_2_ at multiples of the fundamental P_0_ and the narrowness of the peaks as evident from the magnified insets. **E** Modifications of flow rate and therewith of perfusion pump rotation frequency rpm (rotations per minute) are followed by respective changes in P_0_ frequency. Note the weak oscillation P_0_ at 16.6 rpm. **F** The pump rotation (in Hz) multiplied by 8 results in the corresponding P_0_ frequency. Thus the LFP oscillations are directly correlated to the pressure pulsations caused by the 8 rollers of the pump. Created with www.Biorender.com

In our previous publication, we reported on spontaneous slow oscillations in the olfactory bulb (OB) local field potential (LFP) in the NBP (Fig. 2 of Pérez de los Cobos Pallarés et al., 2015). Here we aimed to reveal the origin of these oscillations and uncovered a fast baroreceptive transduction mechanism that couples the pump-induced pressure pulsations to LFP deflections. Based on these observations, we predict a fast influence of heartbeat on neuronal activity in the OB which we confirm with recordings from awake animals. Further, we provide evidence that this transduction is mediated by mechanosensitive ion channels, most likely predominantly by Piezo2 expressed within mitral cells (Wang & Hamill, 2021), which is by now known to be involved in the detection of fast mechanical stimuli throughout the body (reviewed in e.g. Murthy et al., 2017; Szczot et al., 2021). While there are numerous subtypes of peripheral mechanosensory neurons (reviewed in Abraira & Ginty, 2013), mechanosensation by central neurons has rarely been reported so far (e.g. Gaub et al., 2020; Nikolaev et al., 2015).

**Figure 2:**
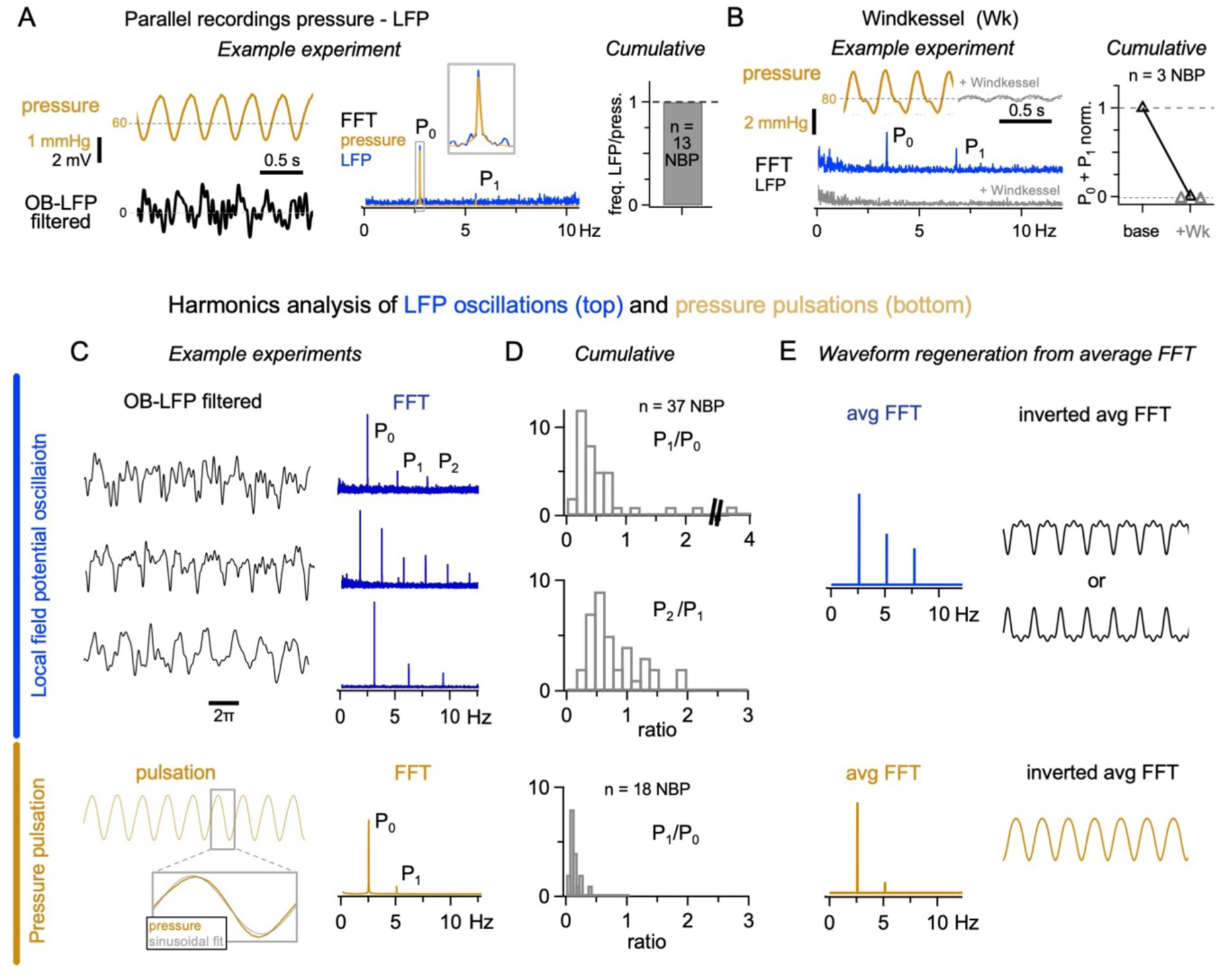
Correlation between OB-LFP oscillations and pressure pulsations **A** Parallel recordings of pressure before the entry point of the aortic cannula and LFP oscillations. Left: Individual example experiment, middle: Overlay of FFT of pressure and OB-LFP recordings. Right: Cumulative data; in all experiments there is a perfect match of P_0_ frequencies for pressure pulsation and slow LFP oscillatory activity. **B** Left: Top: Individual experiment with baseline pressure recording (left) and added Windkessel (right). Bottom: Respective FFTs of the OB-LFP recorded in baseline (blue trace) and with Windkessel (grey trace). Right: Cumulative data of change in (P_0_ + P_1_) amplitude. **C** Example experiments for LFP (top, blue) and pressure (bottom, yellow) recordings with respective FFTs to their right. Time scaled such that one period has the same length for all examples. **D** Cumulative histograms of relative amplitudes of harmonics. No P_2_ was detected in clean pressure recordings (bottom). **E** Left: average power spectra (top LFP, bottom pressure pulsations) based on the average of the cumulative harmonics. Right: inverted FFT of the average spectrum. Top right: Regeneration of the slow LFP in the absence of noise. Note the highly symmetric, denture-like waveforms for the LFP-FFTs (the two renderings are mathematically equivalent) and their match with the examples in C.

The NBP reductionist approach excludes both respiratory patterning (see below) and top-down cortical influences on OB neuronal activity as well as other potential afferents (except for the ethmoidal nerve, see Pérez de Los Cobos Pallarés et al., 2016) and therefore allows to reveal more subtle modulations. Thus, the NBP can serve as an effective tool to investigate so far unknown physiological principles of OB network activity, including the direct physiological effects of heartbeat-induced pressure modulations within the cerebral vascular system on neuronal activity.

Interoception – how the body signals to the brain – plays an important yet so far not fully known role in providing sensory feedback that in turn modulates neuronal activity. For example, respiration is known to strongly pattern neural activity along the olfactory system, not only during odor perception but also spontaneously (e.g. Bolding & Franks, 2018; Carey et al., 2009; Chaput et al., 1992; Gonzalez et al., 2023; Phillips et al., 2012) and even beyond (respiratory rhythm in hippocampus: Lockmann et al., 2016; Nguyen Chi et al., 2016), caused by peripheral mechanosensation of the nasal airflow (Grosmaitre et al., 2007; Iwata et al., 2017). In addition, respiration itself is modulated via mechanosensitive feedback through the stretch receptors in the lung (Hering-Breuer reflex; Kubin et al., 2006; Nonomura et al., 2017), and there are yet more pathways from lung to brain (reviewed in Goheen et al., 2023). Cardiovascular activity so far is known to modulate brain activity mainly via ascending baroreceptive afferents originating from the aortic arch and cardiac tissue (Park & Blanke, 2019, see Discussion). Here we demonstrate another, faster pathway by which arterial pressure pulsations within the brain can directly modulate neuronal activity.

## 2. Methods

### 2.1. The perfused olfactory bulb brainstem preparation (nose-brain preparation, NBP)

All experiments were approved in accordance with the stipulations of the German law governing animal welfare (Tierschutzgesetz). The NBP has been classified as an *in vitro* assay by the local authorities (§ 4 Abs 3 TierSchG; GZ 55.2-1-54- 2532.3-96-13). We have the qualification to perform these experiments as confirmed by AZ DMS-2532-2-74 (Regierung von Unterfranken, Bavaria, Germany). The analyzed experiments are based on a total of n = 145 rats. Different rat strains were used to minimize animal use (the other strains were bred for other ongoing projects in our group) and to improve the significance of results (von Kortzfleisch et al., 2022). Animals were mostly wild-type Wistar rats (70 % of all animals), and to a smaller extent from strains with transgenic expression of GFP variants in either vasopressinergic neurons (VP-eGFP, Wistar, Ueta et al., 2005), GABAergic neurons (VGAT-Venus, Wistar, Uematsu et al., 2008) or dopaminergic neurons (hTH-GFP, Sprague Dawley, Iacovitti et al., 2014). Statistical comparisons between WT and the respective strains during the same experimental phase did not reveal any significant difference regarding the main readout parameter, the oscillatory power amplitude P_0_ of the slow LFP oscillation (see Results, Supplementary Table 1). Thus, these different genetic strains are highly unlikely to interfere with our experimental observations, even more since these transgenic modifications involve only fluorescent labelling of neuronal subtypes that are not essential for the mechanism described here (see Results).

For the NBP, juvenile rats (p15-20 of either sex, not recorded due to the young age of the animals, strains see above) were deeply anesthetized with isoflurane (1-Chloro-2,2,2-trifluoroethyl-difluoromethylether, Abbott, Germany). As described previously in more detail (Dutschmann et al., 2009; Paton, 1996; Pérez de los Cobos Pallarés et al., 2015), once the animal failed to respond to tail pinch, a transection caudal to the diaphragm was performed. Then, the preparation was transferred immediately into an ice-cooled glass chamber filled with artificial cerebrospinal fluid (ACSF; in mM: 1.25 MgSO_4_·7H_2_O, 1.25 KH_2_PO_4_, 3 KCl, 125 NaCl, 25 NaHCO_3_, 2.5 CaCl_2_·2H_2_O, 10 D-glucose). Internal organs including the lungs and heart were dissected. A craniotomy was performed, and the forebrain was removed via gentle suction, leaving the brain stem and the olfactory bulb intact. Afterwards, the left phrenic nerve was isolated and carefully preserved for recording respiratory network activity, while the aorta was separated from the spinal cord and prepared for cannulation. Then, the preparation was transferred to the recording chamber and cannulated through the aorta (Fig. 1A).

Within the recording chamber, the preparation underwent continuous re-oxygenation with carbogenized ACSF (95% O_2_, 5% CO_2_, Linde, Germany) with added sucrose (0.5% to provide oncotic pressure) at a monitored temperature of 30°C (inline heater SH27B, Multichannel Systems, Germany), via a perfusion system using a peristaltic pump (Watson-Marlow 520S-250CA8, UK). A reference electrode was positioned beneath the preparation. Shortly after initiating the perfusion and inserting the cannula into the aorta, spontaneous respiratory movements became evident, which were subsequently suppressed by the introduction of the neuromuscular blocker vecuronium bromide (Sigma; 0.3 μg ml^-1^) into the perfusion system.

### 2.2. Recording phrenic nerve activity and olfactory bulb field potentials, adjustment of perfusion pressure

Glass electrodes were pulled using a pipette puller (Narishige, PC-10, Japan). The size of the tips was widened manually, so that the phrenic nerve ending could be sucked into the electrode to monitor its activity. Recordings were sampled at 1 kHz, amplified, and filtered (DP-311 Differential Amplifier Warner Instruments, USA), digitized via a Powerlab 26T data acquisition device and displayed via LabChart data acquisition software (both AD Instruments, New Zealand). The flow rate of the perfusion system was adjusted to tune the phrenic nerve activity (PNA) pattern according to the known physiological characteristic pattern of eupneic respiratory activity (Dutschmann et al., 2009; Paton, 1996; Stettner et al., 2007); see Fig. 1C for an example). Maintaining a tuned PNA is crucial for our long-term recordings from the olfactory bulb (OB) as a control for the proper oxygenation of the perfused preparation (Pérez de los Cobos Pallarés et al., 2015). The optimal flow rate is influenced by various factors such as the animal’s age, the overall time required for setting up the preparation, and the acquisition and maintenance of an adequate PNA pattern.

Following the tuning and monitoring of the PNA, a glass microelectrode (tip resistance ∼ 2 MΩ) filled with 2 M NaCl) was lowered until touching the dorsal surface of the left OB. Afterwards, a manual manipulator was used to slowly lower the recording microelectrode into the dorsal OB while observing the field potentials of the surrounding cells. The mitral cell layer (MCL) consists of large mitral cell bodies with the same orientation, placed within a single sheet of cells. Mitral cells are known to exhibit substantial spontaneous activity with large amplitude spikes in extracellular recordings (e.g. Chaput et al., 1992). Hence the spontaneous LFP activity recorded from the MCL is robust, especially compared to the external plexiform layer above, which is mostly devoid of cell bodies (Fig. 1B). When robust spontaneous LFP activity from the putative MCL was detected, we recorded long-term LFP as described above for the PNA.

### 2.3 Staining of electrode location

In an initial set of experiments, the localization of the LFP recording site within the MCL was confirmed by injecting Chicago Sky Blue (Sigma-Aldrich, Germany) into the OB via a suction port on the microelectrode pipette holder (Warner Instruments, USA) using a syringe. At the end of each experiment, the OB was isolated from the preparation and fixed immediately by freezing in methylbutane (Sigma-Aldrich, Germany) on dry ice. Next, the bulb was cryosectioned into 40 µm sagittal slices (CM3050 S Cryostat, Leica, Germany) and mounted on microscope slides. The slices were left to dry for 24-48 hours, then a background stain with Ponceau S (0.1% in 5% acetic acid, Sigma-Aldrich, Germany) was performed. Next, the slices were dehydrated in a series of ethanol-dips (Sigma-Aldrich, Germany) with increasing concentration, and cover-slipped using Rotihistol and Histokitt (Carl Roth, Germany). After letting the slides dry at least for 24 hours, the slices were imaged under a light microscope (Leica, Germany).

### 2.4 Pressure recordings in the perfusion system

Pressure measurements were performed to assess the pressure within the perfusion loop, as close as possible to the point where the cannula was inserted into the preparation (see Fig. 1A). An arterial blood pressure transducer (APT300, Hugo Sachs Electronics, Germany) was connected to the input tube next to the entry of the cannula (length 38 mm, inner diameter 0.52 mm, further narrowed tip). The pressure signals were then amplified (CTA342, Hugo Sachs Electronics, Germany), followed by digitization through the PowerLab 26T data acquisition device, also used for PNA and LFP recordings (see above). In a subset of experiments, a Windkessel device was intermittently included in the perfusion system (volume 30 ml, Hugo-Sachs Electronics, Germany), which significantly reduced the amplitude of the pressure fluctuations without changing the mean perfusion pressure. To determine the value of the static pressure at the entry point into the aorta, we performed pressure recordings at a characteristic pump setting (26 rpm/3.5 Hz roller frequency, n = 3 NBPs) before cannulation and after, which yielded a drop in pressure across the cannula alone by ∼ 270 mmHg and a remaining perfusion pressure at ∼ 70 mmHg, i.e. an attenuation to ∼ 20% of the pressure detected before the cannula.

### 2.5 Horizontal and vertical mapping experiments

#### Horizontal mapping

LFP recordings were performed in the dorsal OB from nine different locations, arranged in a grid-like fashion (see also Fig. 5B). The order of the recordings from the different locations varied across the different experiments to avoid possible variances between the physiological state of the preparations during the course of long-term recordings (≤ 2 hrs). In each preparation, we approached the MCL by lowering the recording electrode until the characteristic robust spontaneous MC activity was detected. After finishing the LFP recording from a specific location in the MCL (usually for 360 seconds), the recording electrode was lifted slowly and adjusted to approach a different recording location within the MCL. Dependent on the viability of individual preparations we recorded LFPs from 6-9 locations within a single NBP.

#### Vertical mapping

The recording electrode was incrementally lowered into the dorsal OB, in z-steps of 100 µm from the surface. Several LFP recordings were obtained from each location, on average 8 ± 1. Again, the robust activity of MCs was used to determine the electrode position closest to the MCL (see above). Since the depth of the putative MCL differed between experiments, a total of 4 recordings were analyzed per location and included in the cumulative analysis, a superficial one (glomerular layer - GL), one above the MCL (external plexiform layer - EPL), one at the MCL according to the criterion, and one below (granule cell layer - GCL).

### 2.6 Pharmacology: drug injections

All drugs were pressure injected locally into the OB to avoid potentially confounding effects related to blood-brain-barrier permeability of individual drugs. To do so, a glass pipette (resistance ∼ 5 MΩ) filled with either 30 µM lidocaine (Sigma Aldrich, Germany), 1.25 µM D-GsMTx4 (Sigma-Aldrich, Germany), 10 µM SKF 96365 (Alomone labs, Israel) or ACSF for control, was placed near the recording site. We then recorded neuronal activity of the MCL before and after drug injection.

### 2.7 Nose brain preparation data analysis

#### Fourier and synchronization analysis

For the Fourier analysis of the frequency domains of spontaneous OB oscillations we used IGOR software (WaveMetrics Inc., USA). For comparisons of spectral peak amplitudes between conditions (e.g. baseline vs drug application, different time points at the same location, different strains of animals) of different recording lengths, the power spectra were normalized with respect to the relative duration of the recording(s). For the analysis of treatment effects on spectral peak amplitudes, i.e. on the strength of the oscillatory signal, we often added the amplitude of the first harmonic P_1_ to the fundamental P_0_ to increase the signal to noise ratio.

To analyze the spike phase relative to LFP oscillations, we used custom-written IGOR scripts. This analysis was only performed in experiments with a substantial presence of discernible spontaneous spikes, depending on the electrode impedance, and inherent variance of activity in the tissue; see e.g. Fig. 1C versus Fig. S1D (high versus low spontaneous spike frequency). Briefly, we first took the absolute value of the LFP. Next, we low-pass filtered the original LFP at 10 Hz. The absolute value of this low-pass LFP was then subtracted from the absolute value of the original in order to remove biases due to the ongoing slow oscillation. After subtraction, all negative values were set to 0. Next, spikes were detected using the method from (Drebitz et al., 2019; Quiroga et al., 2004) with the difference that the filtering factor ‘*a*’ in our case ranged from 3 to 4. To prevent over-sampling of a single spike over two adjacent time points, the event with lower amplitude was disregarded. The detected spike times were set to 1 while the remaining time-series was set to 0, hence binarizing the spike signal.

Next, each spike was assigned a phase value with regard to the fundamental slow oscillation. This phase value was derived from the fit of a sinusoidal oscillation (both amplitude and phase) to the low-pass LFP within the analysis interval (60 s). Spikes were sorted into phase histograms (bin width 20°) for each 60 s of the data. The significance of synchronization was assessed with the circular Rayleigh tests and Kuiper tests (available in IGOR) for spike phase histograms obtained for a set of filtering factors *a*, usually 3.0, 3.5 and 4.0. Longer sampling intervals did not necessarily result in higher degrees of synchronization because of the fluctuations both in synchronization and spike rate within individual recordings (see Fig. 3A). We did not optimize interval onset and length for maximal synchrony.

**Figure 3:**
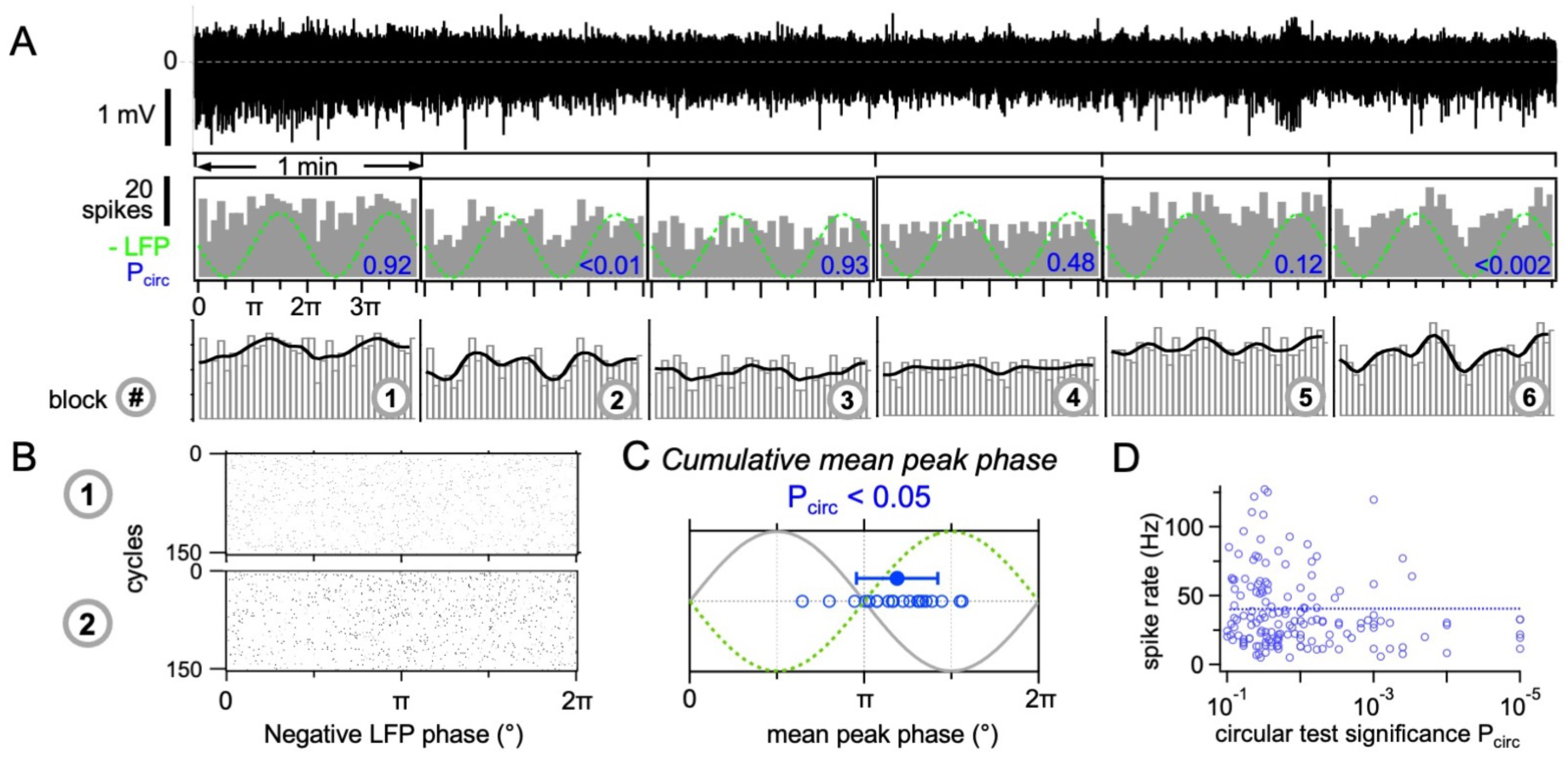
Synchronization of mitral cell spikes to the slow LFP oscillation **A** Example LFP recording (top trace) binned into 1 min-intervals. Middle: Histograms of spike phases within the respective interval (bin width 20°). Green dashed line: inverted P_0_ oscillation at 2.57 Hz (should be correlated to mitral cell membrane potential modulation). Blue numbers: Significance of synchronization (Rayleigh test). Bottom: Smoothed histograms (thick black line). **B** Raster plots of the spikes in block 1 and block 2 shown in A with respect to the negative slow LFP oscillation phase. **C** Mean peak phase of spiking across experiments (averaged over all synchronized intervals with P < 0.05, blue open circles) and average mean peak phase ± S.D. (filled circle, n = 20 NBPs). **D** No correlation between mean spike rate and the significance of synchronization (n = 164 1-minute intervals in 20 experiments). Dashed line: linear fit.

#### Analyses of effects related to changes in pulsation frequency

After the identification of the pressure pulsation as the source of the slow LFP oscillation, we systematically changed pump rates in a set of experiments in various steps of ± 3 rpm relative to the pump rate established in the tuning procedure described above. To test the prediction from Fig. 7A, B – that the adequate stimulus for the baroreceptive transduction mechanism is the positive slope of the pulsation and thus there should be a positive correlation between the maximal slope of the pulsation and the strength of the oscillation - we analyzed the P_0_ amplitudes of the FFTs of similar time windows before and after a change in pump rate and plotted their ratio versus the relative change in pump rate, i.e. the difference between the pump rates before and after divided by the pump rate before (Fig. 7D).

### 2.8 *In vivo* recordings of LFP and heartbeat

All *in vivo* experiments were performed in compliance with German laws and guidelines of the European Union for the use of animals in research (EU Directive 2010/63/EU) and were approved by the local ethical committee (Behörde für Gesundheit und Verbraucherschutz Hamburg, N18/015, N19/021). Adult male and female C57BL/6J mice from the University Medical Center Hamburg-Eppendorf animal facility were housed at a 12/12 h light/dark cycle and had access to water and food ad libitum.

Under isoflurane anesthesia (Abbott, Germany; 2%), mice were implanted with two plastic bars for head fixation. The skull above the OB (coordinates from bregma: 5.0 mm anterior, 0.8 mm lateral) was exposed and electrode insertion sites were marked on the skull. A silver wire was implanted above the cerebellum as reference electrode. After implantation, mice received buprenorphine (0.1 mg/kg) and recovered in their home cage for at least 5 days. Afterwards, mice were accustomed to handling and head fixation.

For the recording, a small craniotomy (<0.5 mm in diameter) on the marked site above the OB was performed under brief isoflurane anesthesia (1%), mice were loosely fitted into a plastic tube, head fixed, a sterile syringe needle was inserted subcutaneously in the back to record the electrocardiogram (ECG), and a silicon probe was inserted into the ventral OB (16-channel, 50 μm spacing, Neuronexus, USA, ∼ 2.5 mm from the dura). The ventral OB was recorded *in vivo* to reduce any potential impact of brain movement due to the craniotomy. The reversal of the polarity of the respiration related LFP served as orientation to position the electrode spanning the MCL. Respiration was recorded using a pressure sensor (Honeywell, AWM3300V, Germany) in front of the nose as previously described (Bitzenhofer et al., 2022).

The mouse was allowed to recover from isoflurane anesthesia for ∼ 45 min before recordings were started. Extracellular signals were band-pass filtered (0.1 - 9000 Hz) and digitized (32 kHz) with a multichannel extracellular amplifier (Digital Lynx SX, Neuralynx, USA). ECG and respiration (band-pass filtered 0.5 - 500 Hz) were recorded with the same amplifier. For some recordings, probes were coated in DiI to verify recording locations post hoc. Some mice were used for up to 3 recording sessions over several days to reduce the animal numbers (5, 4, 2 mice with 1, 2, 3 sessions, respectively) by covering the craniotomy with silicon sealant after the first recording. Distinct probe tracks in these mice make it unlikely that the same neurons were recorded across sessions.

For recordings with naris occlusion, the nostril ipsilateral to the recording site was sealed reversibly with silicon elastomer.

### 2.9 Analysis of *in vivo* data

Data were analyzed with open source and custom-written algorithms in the MATLAB and Python environment.

#### Inhalation detection

The respiration signal recorded with a pressure sensor in front of the nose was filtered at 1 - 30 Hz and smoothed with a 40 ms sliding window. Potential inhalation onsets were detected as negative deflection peaks with a peak prominence above 1 standard deviation on the difference between adjacent samples (pressure change). Snippets of 200 ms centered around the inhalation onset were clustered into 25 clusters using k-means clustering. The clusters were manually inspected and labelled as normal inhalation, sniffing, and noise. Only normal inhalations were considered for inhalation alignment of single unit activity.

#### Heartbeat detection

The ECG signal was filtered at 10 - 300 Hz. Potential heartbeats were detected as negative and positive deflection peaks with a peak prominence above 3 standard deviations. Snippets of 15 ms centered around the heartbeat onset were clustered into 25 clusters using k-means clustering. The clusters were manually inspected and labelled as positive ECG peak (R-peak), negative ECG peak (S-peak), and noise. S-peaks were primarily considered for heartbeat alignment of single unit activity. R-peaks were used in segments of the signal with unreliable S-peak detection.

#### Reversal channel detection

Linear 16-channel probes with 50 μm distance of recording sites were used for LFP recordings in the ventral OB (Fig. 4A, B). Electrodes were positioned according to the respiratory LFP which reverses in polarity close to the mitral cell layer (Fourcaud-Trocmé et al., 2014). For group analysis, recording sites from different recordings were aligned to the reversal of the respiratory LFP. Extracellular recording signals were filtered at 1 - 12 Hz. Snippets of 1000 ms centered around 1000 inhalation timestamps were averaged per recording and the absolute peak value 0 - 200 ms after inhalation was calculated for each channel. The channel with the lowest value was considered the reversal channel.

**Figure 4:**
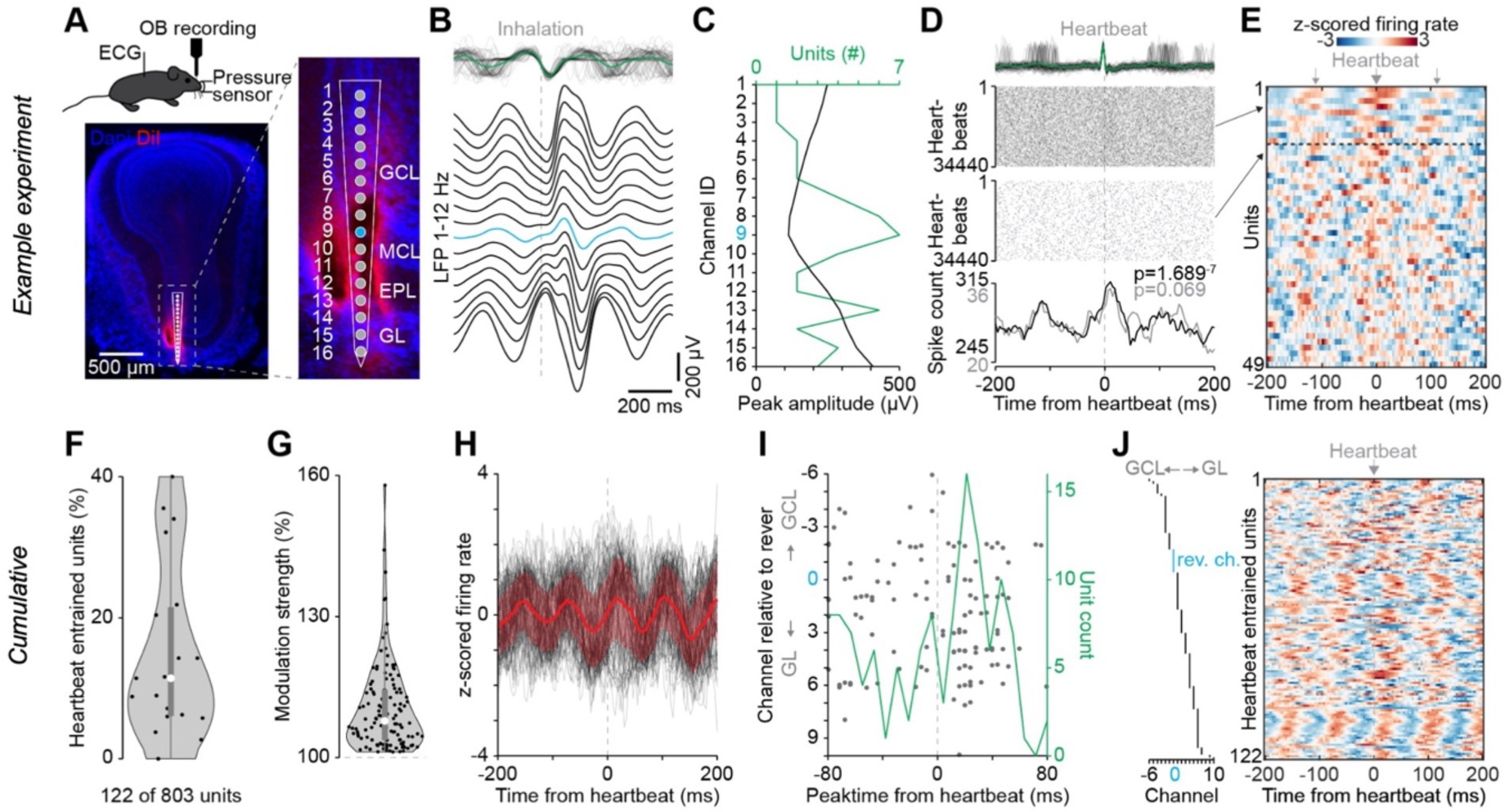
Heartbeat entrainment of bulbar neurons *in vivo* **A** Left: Scheme of the *in vivo* recording setup and below example coronal section from one mouse showing silicon probe track labelled with DiI in OB. Right: Reconstruction of the position of the 16 recording channels in the ventral OB. Blue signifies the reversal channel. **B** Example LFP (1-12 Hz) aligned and averaged for 1000 inhalations for all channels. Pressure signal measuring respiration is shown at the top. **C** Example of detection of the reversal channel (blue) based on the respiration-related LFP activity and its relationship to the number of detected units. **D** Raster plots (top) and peri-event time histograms for two example units relative to heartbeat. ECG signal is shown at the top. **E** Color-coded z-scored firing rate for each unit in an example recording session relative to heartbeat. Each row shows the activity of a single unit sorted by ascending p-value from top to bottom. Single units above the dashed line are significantly modulated by heartbeat (Rayleigh circular test, p < 0.05). **F** Percent of heartbeat entrained units for 19 recording sessions from 11 mice. **G** Modulation strength of heartbeat entrained units. **H** Z-scored firing rate of significantly heartbeat entrained units aligned to heartbeat. Red is the average of all significant units. **I** Distribution of peak firing rate time points of heartbeat entrained units versus channel number relative to the reversal channel (blue 0) and the corresponding histogram (green). **J** Z-scored firing rate of significantly entrained units aligned to heartbeat (right panel), arranged in the order of their location with respect to the reversal channel (blue, left column).

#### Single unit activity

Single unit activity was detected and sorted using Klusta program (Rossant et al., 2016), followed by manual curation in phy (https://github.com/cortex-lab/phy). Peri-event time histograms were calculated for each unit around the inhalation timestamps and heartbeat timestamps. Entrainment of single units to respiration and heartbeat was tested using the Rayleigh circular test (p < 0.05 considered significant). Modulation strength was calculated from peri-event time histograms as the maximum divided by the mean firing rate.

## 3. Results

### 3.1 Slow LFP oscillations in the NBP are related to pressure pulsations induced by the peristaltic pump

The present study aimed to further investigate the origins of spontaneous oscillatory activity in the *in situ* perfused OB (Pérez de los Cobos Pallarés et al., 2015). Oscillations within a frequency range of 1.5 – 4 Hz (Fig. 1C, Fig. S1A) were recorded within or very close to the mitral cell layer (MCL; Fig. 1B; see Methods) and observed in the majority of NBP preparations that showed regular PNA bursting (burst frequency at 14.4 ± 5.3 bursts/min, n = 38 analyzed experiments). Note that in contrast to previous reports on *in vivo* studies (e.g. Buonviso et al., 2006; Rojas-Líbano et al., 2014) MCL oscillations *in situ* are not coupled to the respiratory rhythm due to the absence of nasal airflow and the severed ascending fiber tracts that could have provided inputs from respiratory brainstem networks. Thus, the *in situ* perfused OB is devoid of any respiratory modulation, even though rhythmic PNA is generated by the brainstem and serves as an important readout for the overall viability of the perfused NBP (Pérez de Los Cobos Pallares et al., 2016, Fig. 1C, S1C, D).

The Fourier analysis further revealed harmonics of this fundamental rhythm (P_0_) in 53 out of 80 analyzed experiments, with on average 2.0 ± 1.0 detectable harmonic peaks (P_1_, P_2_ etc., Fig. 1D). The individual frequency peaks were very narrowly tuned (Fig. 1D, S1A; P_0_ average full width half maximum FWHM = 4.9 ± 3.3 mHz, n = 55). We did not notice this narrowness in our original publication of the NBP because of the filter setting used for the Fast Fourier Transform (FFT) (Pérez de los Cobos Pallarés et al., 2015; see e.g. their Fig. 2D). Since slow LFP oscillations such as theta oscillations in the hippocampus, and also respiration-related oscillations in the OB show far larger spectral widths (e.g. Buzsáki, 2002; Lockmann et al., 2016; Karalis & Sirota, 2022), we raised the question whether the slow oscillations might be an electrical or mechanical artefact related to the perfusion pump. However, changing the placement of the reference electrode did not influence the oscillatory signal (see Fig. S1B). Moreover, the peristaltic perfusion pump was operated well below 1 Hz at 15 - 25 rpm (rotations per min) to provide flow rates for optimal perfusion.

Further investigation into the origins of a potential mechanical artefact finally revealed that the periodic LFP signal was entrained to the pulsations caused by the 8 rollers of the peristaltic pump and thus occurred at a frequency 8 times the pump rate (Fig. 1E, F). Importantly, the setting of the pump rate was determined by the tuning process that was used to achieve physiologic PNA (see Methods), thus the fundamental slow oscillation frequency appeared to be biologically distributed across NBPs (Fig. S1A).

Nevertheless, the observed LFP oscillations must be of neuronal origin since they were associated with the viability of the *in situ* perfused olfactory bulb. In n = 7 experiments we investigated the impact of hypoxia on slow LFP oscillations. We recorded baseline LFP activity for at least 10 mins before we lowered pump rate to 3 rpm to evoke progressive hypoxia for 10 mins until we re-established the initial pump rate. In all experiments (Fig. S1C), the rhythmic PNA displayed the characteristic hypoxic response pattern (Powell et al., 1998) of augmentation followed by depression and finally cessation of bursting activity. During the stage of hypoxic synaptic depression (see Richter et al., 1999) PNA disappeared, and the slow LFP oscillation also either disappeared entirely or its power decreased substantially (a systematic rundown in P_0_ and P_1_ can be excluded, see stationarity analysis in Fig. 5A and Fig. S5B). Similar observations were made in a subset of long-lasting experiments in which the PNA bursting eventually stopped spontaneously due to a rundown of the preparation (n = 6 NBPs, Fig. S1D).

**Figure 5:**
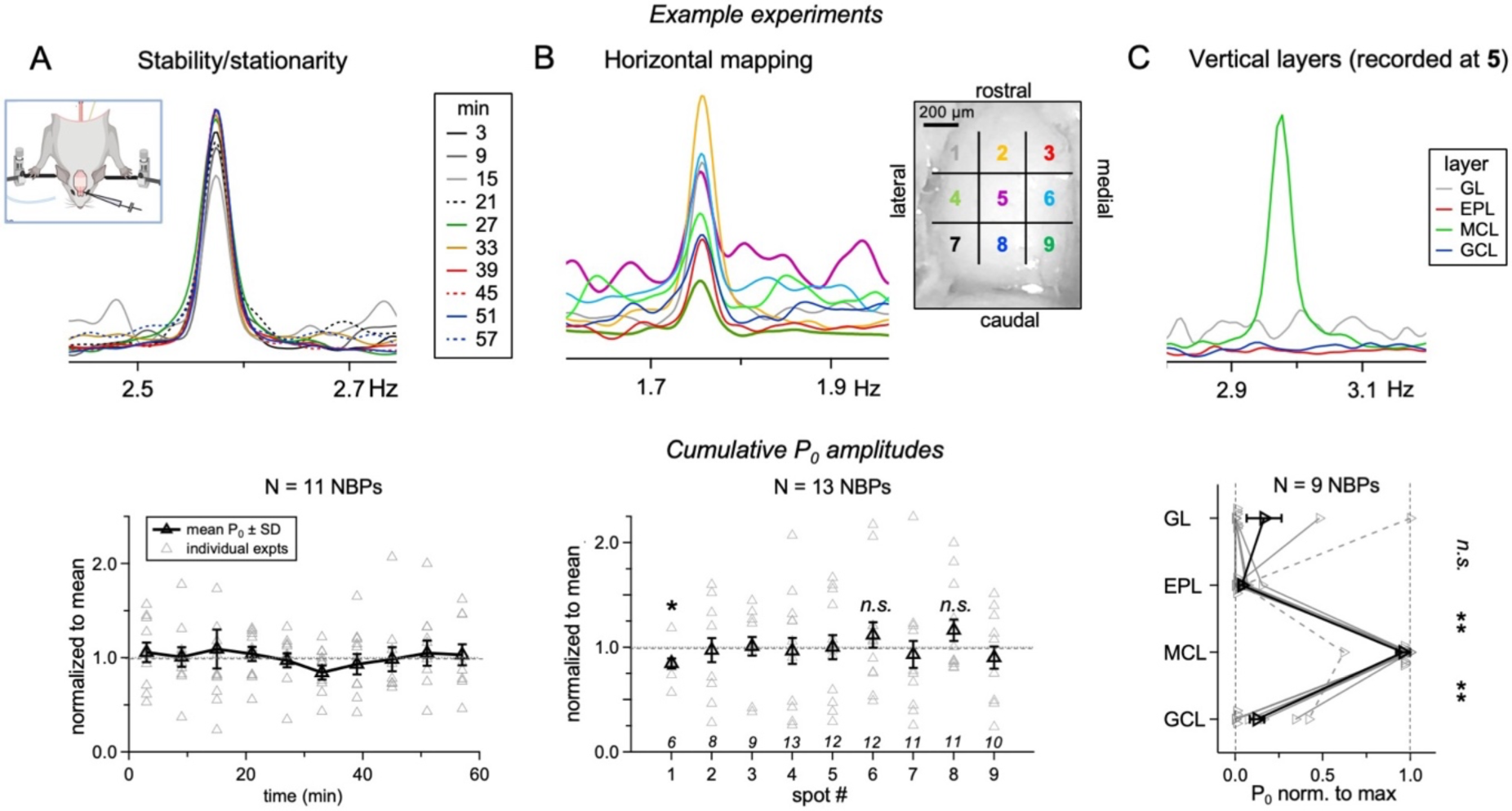
Slow oscillations over time, horizontal location and the layers within the OB For all types of experiment, a representative example of the FFT set of an experiment is shown in the top panel and cumulative data at the bottom. **A** Recording for up to 60 minutes, data binned into 6 min-intervals and normalized to the mean P_0_ amplitude within experiments. Huge variability within experiments, but no systematic rundown. **B** Recording from up to 9 locations across the OB surface within the same NBP. Normalized to the mean P_0_ amplitude within experiments. Again, substantial variability, larger than for temporal stability, but no systematic deviation from the mean except for location **1** with a slightly smaller average. Small numbers above bottom axis indicate number of data points recorded at the respective location. **C** Recording from the same location at the central dorsal OB (**5**) but at different vertical depths, spaced at 100 µm. Since the thickness of the OB layers and thus layer localization are variable and cannot be determined by depth alone, representative recordings for each layer (1 each) were selected. The mean P_0_ recorded from the vicinity of the mitral cell layer (MCL) is significantly larger than those recorded from either putative external plexiform or granule cell layer (EPL, GCL). In a small subset of experiments (n = 2) there was also a substantial contribution in the glomerular layer (GL).

### 3.2 Slow LFP oscillations are a baroreceptive response and not driven by synaptic input

To directly establish the correlation between the LFP oscillations and the rhythmic pulsations caused by the roller pump, we next recorded OB LFPs and pressure simultaneously (see Methods, Fig. 1A). Perfusion pressure recordings displayed pulsations with an amplitude of ∼ 2 - 4 ΔmmHg on top of the static pressure established by the pump (∼ 40 - 80 mmHg at entry point to aorta, Fig. 2A, B, see Methods). The FFT of the pulsatile signal revealed exactly the same fundamental frequency as the slow LFP oscillation (Fig. 2A; mean ratio of frequencies 1.0 ± 0.0, n = 13 NBPs). As shown in Fig. 2B, a reduction of the pulsation amplitude to ∼25% of the baseline via a switched inclusion of a Windkessel device in the perfusion system (see Methods) resulted in complete cessation of the slow LFP oscillation (n = 3 NBPs), providing further causal evidence for a fast baroreceptive transduction mechanism. These data from the *in situ* NBP imply that the slow LFP oscillations in the olfactory bulb are caused by baroreceptive coupling, which can be further investigated via the analysis of harmonics.

Harmonics reflect the deviation of an oscillatory signal from a purely sinusoidal waveform and are to be expected for neuronal network oscillations because of the asymmetry of postsynaptic potentials’ time courses (illustrated e.g. in Scheffer-Teixeira & Tort, 2016, their Fig. 4). The top panels of Fig. 2C show a set of characteristic slow LFP waveforms and their corresponding FFT spectra from different NBPs. Fig. 2D top illustrates the cumulative distribution of harmonics relative to the fundamental frequency for a set of experiments (n = 37) with the power of the first harmonic peak P_1_ at, on average, 56 ± 67 % of the fundamental P_0_, and the power of the second harmonic peak P_2_ at 77 ± 42 % of P_1_. Fig. 2E top shows the average FFT based on average P_1_/P_0_ and P_2_/P_1_ ratios and its inverse, reconstructing the slow oscillatory component of the LFP. As evident from the bottom panels of Fig. 2C, D, E, the recorded pressure pulsation was mostly sinusoidal in nature with a minor harmonic addition (P_1_ = 10 ± 8 % of P_0_, n = 18 experiments with pressure recordings).

The LFP reconstruction in Fig. 2E shows a remarkable mirror symmetry, which can also be observed in the filtered data examples shown in Fig. 2C. Thus, the waveform analysis excludes synaptic origins of the observed LFP modulation since coordinated action potentials and postsynaptic potentials in response to synaptic inputs would rather predict an asymmetric pattern (e.g. Scheffer-Teixeira & Tort, 2016; reviewed in Cole & Voytek, 2017). These observations, together with the narrowness of the FFT peaks, imply that a direct baroreceptive transduction mechanism is the most likely explanation for the generation of slow LFP oscillations, rather than a mechanism that involves neuronal network activity. Next, we hypothesized that if the pressure pulsations did indeed induce rhythmic electrical signals in the mitral cells, their spontaneous spiking activity should also be modulated by the pulsation-induced oscillatory LFP.

### 3.3. Spontaneous MC activity is synchronized to the oscillatory LFP

It has been repeatedly demonstrated *in vivo* that spontaneous mitral and tufted cell activity is often modulated by the respiration-related theta rhythm 8_RR_ (e.g. Briffaud et al., 2012; Fukunaga et al., 2012); thus, we tested whether such a synchronization would also occur for the pressure pulsation-induced LFP oscillation in the absence of any respiration-related sensory feedback. In n = 21 experiments we obtained LFP recordings that allowed for the detection of multi-unit activity in the MCL (Fig. 3A, B). Since multi-unit activity fluctuated substantially over time, we analyzed synchronization within time blocks of 1 min, in a total of 426 blocks with a mean spike rate of 45 ± 35 Hz (mean block number 20 ± 20 per experiment, range 6 - 60). All n = 21 experiments contained blocks with significant modulation as determined by circular statistical tests (see Methods; in total n = 141 significant blocks with a mean spike rate of 40 ± 32 Hz, no correlation between spike rate and significance: r = 0.02, P = 0.37, Fig. 3D).

As to the preferred phase of synchronization, the phase maximum (averaged over significant blocks within experiments, n = 21) was located on average 209 ± 46° with respect to the fundamental frequency of the slow LFP oscillation. The spontaneous spiking significantly synchronized with the 2^nd^ half of the rising phase of the inverted LFP oscillation at the fundamental frequency (Fig. 3C, P_binomi_ = 0.0000013 for interval π - 3π/2, P_binomi_: cumulative binomial probability for the observed distribution of data points). This entrainment of mitral cell spiking to the rising phase of the inverted LFP is to be expected, if the inverted LFP oscillation corresponds to the modulation of the intracellular mitral cell membrane potential – in line with a direct involvement of mitral cells in the generation of the oscillatory LFP.

In summary, in the rat NBP spontaneous mitral cell activity is in part and intermittently modulated by pressure pulsations induced by the perfusion system, the frequencies of which fall within the physiologically relevant frequency regimes of both respiratory rhythm and heartbeat frequency (see Discussion). Such synchronization argues in favor of a biological, baroreceptive mechanism that directly modulates mitral cell excitability, irrespective of any modulatory afferent synaptic inputs.

### 3.4 A subset of olfactory bulb neurons is entrained by heartbeat *in vivo*

Based on the observed synchronization of mitral cells with the pulsation-induced slow LFP oscillations *in situ*, we hypothesized that pulsations of intracranial pressure should modulate neuronal activity in the OB *in vivo.* Indeed, heartbeat related pressure changes on the order of 5 - 10 mmHg in humans (Wagshul et al. 2011) and rodents (Eftekhari et al. 2020) are at a similar range compared to the pulsatile pressure changes introduced by the peristaltic pump in the NBP (see above). To test this hypothesis, we simultaneously recorded the ECG, nasal airflow, and OB activity in awake, head fixed mice (n = 19). Linear silicon probes with 16 recording sites were used to record neuronal activity in the ventral OB (rather than dorsal for increased mechanical stability; Fig. 4A). For analysis, channels from different recordings were aligned to the reversal of the respiration-related activity (Fig. 4B, C). We observed a modulation of the firing rate relative to heartbeats in a subset of neurons as shown in the peri-event time histograms of single units (Fig. 4D, E). In total, 122 of the 803 units recorded in 19 recording sessions in n = 11 mice were significantly entrained by the heartbeat (Fig. 4F). While the activity of OB neurons was modulated by heartbeat, the ensuing variation of their firing rate was mild. We calculated the modulation strength (peak firing rate / mean firing rate) to quantify the effect size and found that the firing rate peaks of significantly entrained units were about 10% above their average firing rate (Fig. 4G). The firing rate of the significantly entrained units predominantly peaked ∼ 20 ms after the heartbeat (Fig. 4H-J). With a pulse wave velocity of ∼ 3 m/s (Hartley et al., 1997) and a heart to head distance of ∼3 cm in mice, this timing is consistent with a heartbeat-related pressure modulation of OB activity and excludes a modulation via multi-synaptic inputs arising from aortic or carotid baroreceptors (see also Discussion).

Both, the number of significantly entrained units and the modulation strength were higher for nasal airflow (Fig. S4A, B, C), the main driver of activity in the OB, peaking at ∼ 150 ms after the onset of inhalation (Fig. S4D, E). While they operate at different timescales, respiration and heartbeat are not independent (Dick et al., 2014) and 119 of the 122 units that were significantly modulated by heartbeat were also modulated by respiration. To further exclude that heartbeat entrainment is a side effect of cardiorespiratory coupling we performed additional recordings of OB activity, ECG, and respiration in n = 6 mice with and without occlusion of the naris ipsilateral to the OB recording site (Fig. S4F-J). While naris occlusion significantly reduced the number of significantly entrained units and the modulation strength relative to nasal airflow (Fig. S4G, H), the influence of heartbeat on OB spiking activity was unchanged (Fig. S4I, J). In summary our data provide *in vivo* evidence that the activity of a subset of neurons in the OB is modulated by heartbeat-induced pulsations.

### 3.5. Slow LFP oscillations in the NBP are robust and localized to the mitral cell layer

Next, we aimed to investigate the molecular underpinning of the baroreceptive mechanism of OB oscillations using pharmacology *in situ*. In a set of control experiments we first investigated the experimental robustness of the OB oscillation. To do so, LFPs were recorded from the same location and with the same pump settings for over 60 minutes in n = 11 NBPs. These recordings show that the oscillation remained unchanged within 10 consecutive blocks of 6 min LFP recordings (Fig. 5A). While P_0_ peak power was variable across intervals (mean CV 0.38 ± 0.21, range 0.17 – 0.81), no systematic overall variation was detected (no significant difference of any interval data set from the distribution within the first interval, Wilcoxon test). Only the overall LFP activity in terms of the LFP standard deviation, which was also highly variable (same set of experiments, mean CV 0.20 ± 0.15, range 0.02 – 0.51), declined over time (linear fit slope = - 0.5 %/min, Fig. S5A), whereas there was no rundown in the baroreceptive response to pulsatile stimulation at 2 - 4 Hz.

In other sets of experiments, we investigated whether the pulsation-induced oscillation displays topography within the dorsal OB. Thus, we recorded sequentially from up to 9 locations arranged in a grid (Fig. 5B) on the dorsal surface of the OB (5 ± 2 locations/preparation, n = 17 NBPs). Similar to the temporal stability recordings above, variability in power P_0_ was high, with a mean CV of 0.60 ± 0.31, but the observed local deviations from the normalized mean across the individual experiments were not significant, except for the antero-lateral grid location **1**. Here the increased anatomical curvature of the bulbar layers (e.g. Herreras, 2016) might have contributed to significant reduction of P_0_ power. However, our mapping analysis showed differences for the localization of the slow LFP oscillations across the glomerular, external plexiform, mitral and granule cell layers, as recorded from the central dorsal olfactory bulb (see Methods, n = 9 NBPs). Here, P_0_ of the slow oscillation was localized predominantly in the MCL (Fig. 5C).

In addition, we evaluated the spatiotemporal stability of harmonics (Fig. S5B, C). The same control experiments revealed no significant fluctuation in neither the occurrence of harmonic peaks P_1_ and P_2_ nor P_1_/P_0_ power ratio over time (Fig. S5B, n = 5 experiments) or across mapped locations (Fig. S5C, n = 13 experiments).

In conclusion, these control mapping and long-term recording experiments suggest that pressure pulsation-induced oscillations are robust across the dorsal OB and restricted to the vicinity of the MCL. Conversely, tufted cells and granule cells and others apart from mitral cells are unlikely to contribute to such oscillations.

### 3.6. Pharmacology: Baroreceptive LFP oscillations are independent of network activity but require mechanosensitive ion channels

Because of the symmetric waveform of the slow oscillation and its independence of mitral cell spiking activity we hypothesized that neuronal network activity (both synaptic transmission and action potential firing) would not be involved in the putative transduction mechanism for pressure pulsation.

An initial set of control experiments using local ACSF microinjection showed no significant changes in fundamental P_0_ power, spike count or LFP standard deviation (n = 6 NBPs, Fig. 6A, Table S6). Subsequently, we pharmacologically blocked voltage-gated sodium (Na_v_) channels via local lidocaine injection (n = 8 NBPs, Fig. 6B). Lidocaine caused significant reductions (Mann-Whitney-test P < 0.01) in LFP standard deviation (63 ± 33% of baseline) and spike count (25 ± 33% of baseline), while the mean oscillation power P_0_ + P_1_ was not significantly changed (130 ± 88% of baseline). The lidocaine experiments further support our hypothesis that the transduction of pressure oscillations does not require spiking activity of mitral cells or other neurons.

**Figure 6:**
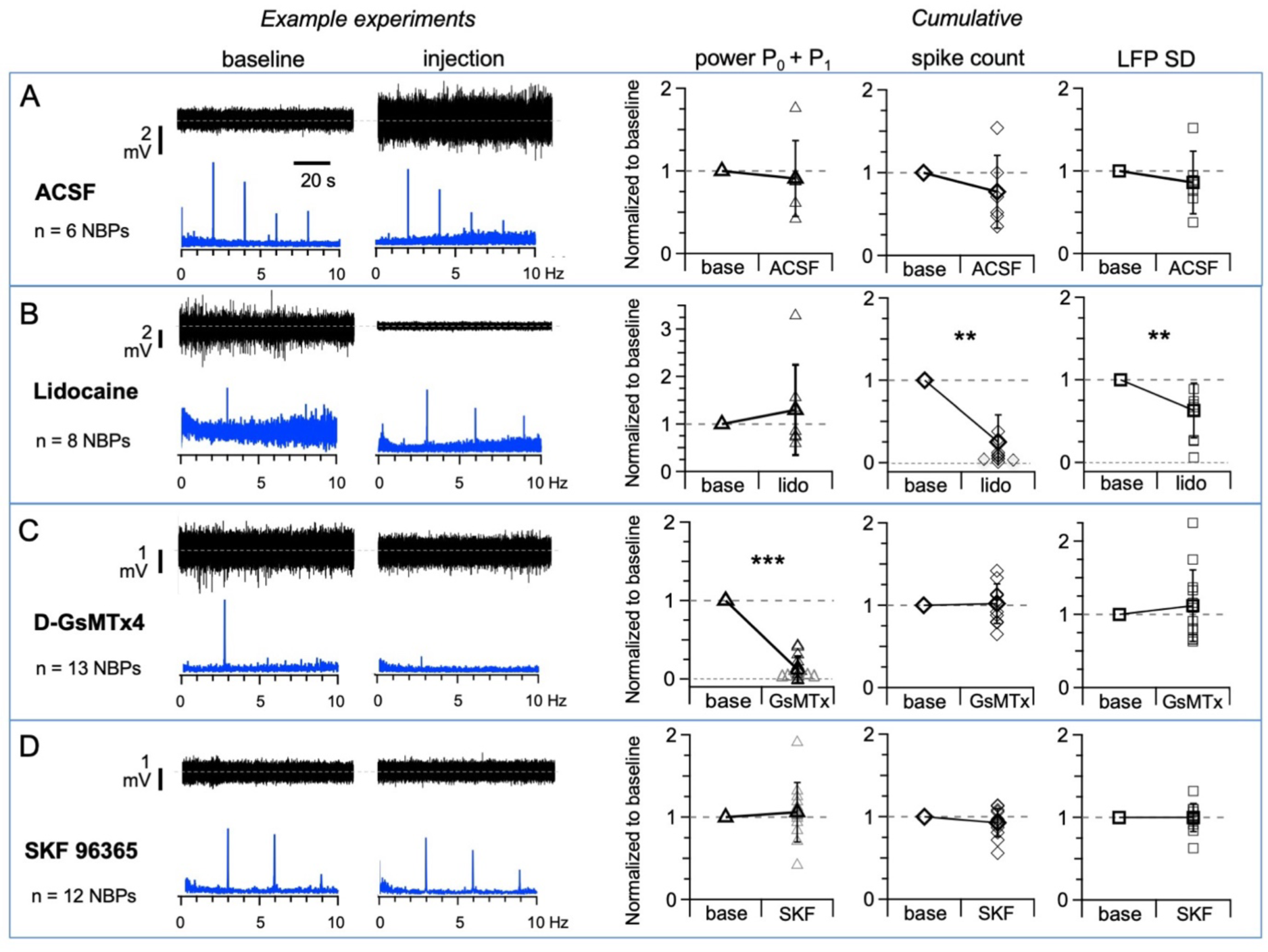
Slow LFP oscillation abolished by blockade of mechanosensitive ion channels but not by blockade of neuronal spiking For all conditions, baseline and post injection data from an example experiment are shown on the left and cumulative data on comparison of parameters oscillatory power, spike count and LFP activity during baseline and post injections normalized to baseline are shown on the right. The FFT data show no scaling since they were normalized to adjust for different recording times in both conditions. **A** Control for injection technique: Injection of ACSF close to the recording site. No significant changes were observed. **B** Injection of lidocaine (30 µM, Na_v_ channel blocker) resulted in significant reductions of spike count and LFP activity but did not influence oscillatory power. **C** Injection of D-GsMTx4 (1.25 µM, blocker of cationic mechanosensitive channels) resulted in significant reduction of the oscillatory power, while there was no effect on spike count and LFP activity. **D** Injection of SKF 96365 (10 µM, unspecific TRP channel blocker) did not influence any parameter.

Since the baroreceptive mechanism appears to directly modulate cellular excitability without involving synaptic transmission, we tested the hypothesis that pressure pulsations might be transduced via excitatory/cationic mechanosensitive ion channels. Interestingly, in the OB Piezo2 expression has recently been localized specifically to a subset of mouse mitral cells via both immunohistochemical and RNAseq approaches (Wang & Hamill, 2021; Zeppilli et al., 2021). Only a few pharmacological tools are available to selectively block mechanosensitive ion channels. GsMTx4 (the spider toxin *Grammastola spatulata* mechanotoxin 4, (Suchyna, 2017; Suchyna et al., 2000) is known to interfere with the gating of various cationic mechanosensitive ion channels including Piezo1, Piezo2 and TRPC1 and TRPC6 channels at similar concentrations (Alcaino et al., 2017; Gnanasambandam et al., 2017). In line with our working hypothesis local microinjection of D-GsMTx4 (1.25 µM, n = 13 NBPs) resulted in either a substantial decrease or complete block of slow oscillations in all experiments (to 13 ± 11 % of baseline, Fig. 6C, P < 0.001 Wilcoxon-Test), while no significant changes in spike count or LFP standard deviation (94 ± 24 % and 125 ± 59 % of baseline, respectively) were observed. While antagonistic effects of D- GsMTx4 on Na_v_ and K_v_ channels have been reported (IC_50_ in 10 µM range, Redaelli et al., 2010, vs 1.25 µM used here) we conclude that the toxin selectively reduced the pressure pulsation-driven LFP oscillations since overall network activity was preserved.

Next, we injected 10 µM SKF 96365, a broad-band blocker of the TRPC channel family (transient receptor potential type C channels; e.g. Grotle et al., 2021), of which several members are also expressed in mitral cells (Dong et al., 2012; TRPC1, 3 and 5 according to the data base associated with Zeppilli et al., 2021). SKF 36365 did not cause any effect on oscillations, spike count or LFP standard deviation alike (n = 12 NBPs, 106 ± 36%, 93 ± 17% and 100 ± 16%, respectively, Fig. 6D), thus TRPC channels are unlikely to be involved in the generation of pulsation-induced slow oscillations.

### 3.7. Piezo2 channel properties predict symmetric LFP and positive correlation between pulsation slope and LFP oscillation power

The above pharmacological experiments together with the baroreceptive LFP waveform suggest that synaptic transmission is not involved and that Piezo channels could be a possible substrate for the transduction of pressure pulsations in mitral cell membranes and thereby enable direct modulation of mitral cell excitability. Piezo channels have been shown to provide the basis for fast, differential mechanosensing in the skin and other body tissue types such as the lung and the gastrointestinal tract (e.g. Alcaino et al., 2017; Min et al., 2021; Nonomura et al., 2017) but also in baroreceptive neurons in the nodose ganglia where they actually contribute to mediation of the baroreflex (Zeng et al., 2018). While the gating mechanism of these channels is not yet fully understood and different types of mechanical stimulation might drive different types of gating (reviewed in Young et al., 2022), many characteristics of Piezo channel operation are known by now.

Gating of Piezo channels is very fast (activation << 1ms) and involves an inactivated state (Piezo1 inactivation < 20 ms, Piezo2 inactivation < 3 ms at resting membrane potentials; e.g. Coste et al., 2010; Moroni et al., 2018, reviewed in Young et al., 2022). Thus Piezo-mediated baroreceptive transduction is differential, i.e. sensitive to changes in pressure and not to static pressure, and can directly follow fast pressure pulsations, as e.g. during tactile exploration of a textured surface by somatosensory mechanoreceptive neurons (Lewis et al., 2017).

What causes the observed transformation of the pressure pulsation waveform into the LFP waveform (Fig. 2E)? In spite of the structural similarity of the two Piezo channels, Piezo2 is likely to be substantially more responsive to increases in pressure than to decreases, whereas Piezo1 and probably most other baroreceptive channels are sensitive to both types of stimuli (e.g. Moroni et al., 2018; Shin et al., 2019; reviewed in Young et al., 2022). Accordingly, Piezo2’s open probability P_open_ should be highest during the maximum upwards slope of the pulsation (Fig. 7A). Due to the fast inactivation of the channel compared to the time scale of the pulsation (< 10 ms vs 100 - 200 ms), the resulting current should closely follow P_open_, resulting in rhythmic depolarization at the same frequency as the pulsation and yield a mostly symmetric signal, since the local LFP is generated by the spatial summation of ionic currents. Conversely, Piezo1 or other mechanosensitive channels would be expected to be gated at double the pulsation frequency and in addition should produce asymmetric currents because of the extended inactivation compared to Piezo2 – as actually observed in Piezo1 currents in response to sinusoidal hypobaric stimuli (at 1 - 5 Hz, Lewis et al., 2017). Indeed, the previously reconstructed shape of the slow LFP oscillation can be explained very well by the proposed Piezo2 response pattern (Fig. 7A bottom). Since we had reconstructed the slow LFP before investigating its molecular mechanism (Fig. 2E), there was no fitting in hindsight.

**Figure 7:**
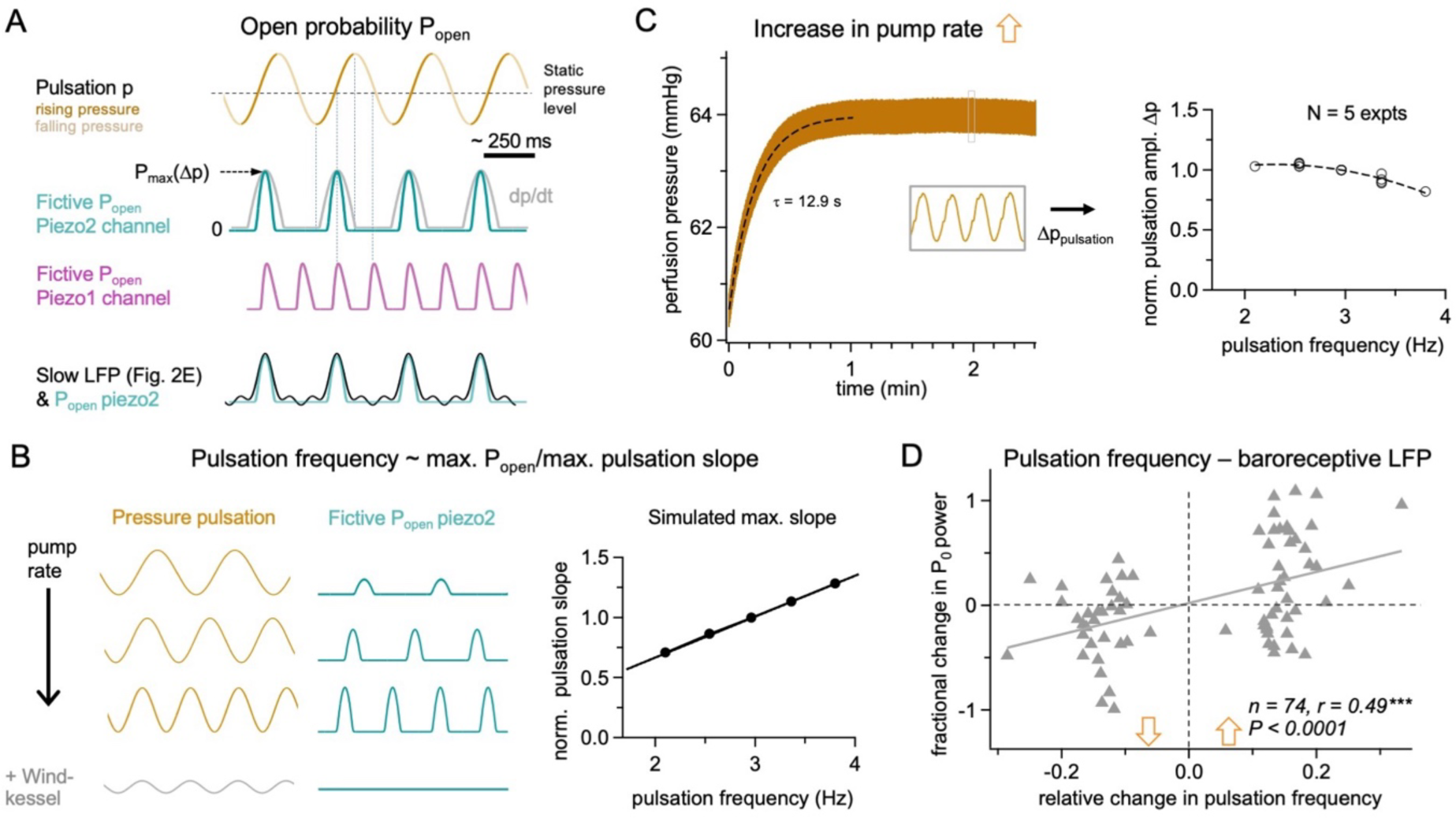
Biophysics of Piezos – predictions and data tests. **A** Fictive gating of Piezo channels in response to sinusoidal pressure pulsations at ∼2 Hz according to known gating properties. Top trace: Pressure pulsation stimulus, phases of increasing pressure labelled in dark yellow. Next: Open probability P_open_ of Piezo2 (green trace) should be highest during maximal positive slope of pressure (differential sensor, rectifying). Since the change in slope might not be sufficient for gating during the onset and end of the positive pressure increases, the actual opening is likely to be restricted to an interval shorter than the first derivative of the rising pressure intervals (green trace vs grey trace). Due to the fast inactivation (<3 ms), P_open_ should be symmetric around the maximal positive slope. Next: Piezo1 or other mechanosensitive channels are likely to be gated during the higher slope sections of both increasing and decreasing phases of pressure levels (although the maximal P_open_ might differ between the two types of gating; altogether differential sensor, non-rectifying). Due to the longer inactivation compared to activation (or lack of inactivation), gating should be less symmetric (purple trace). Lowest traces: Overlay of the fictive P_open_ of Piezo2 (green trace) with the slow LFP waveform recreated from the average harmonics (black trace, Fig. 2E). **B** With increasing pump rate, pulsations will speed up (left side) and thus maximal slope and therewith maximal P_open_ (middle) should also increase. Right side: Calculated normalized maximal slope (first derivative) of sinusoidal pulsation versus pulsation frequency. The addition of the Windkessel device (Fig. 2B, bottom) reduces pulsation amplitude and therewith maximal slope; in our experiments, there was no more slow LFP oscillation in the Windkessel condition. **C** Left: Recorded characteristic example of pressure change directly after an increase in pump rate by 3.2 rpm at time point 0. The static pressure level assumes its new value with a time constant of ∼10 s (dashed black line; top graph on the right shows relation between static pressure level and pump rate, filled black circles from n = 5 experiments and linear fit). Pressure pulsations are evident from the magnified inset. Right: Pressure pulsation amplitudes do not increase with increased pump rate (right, open black circles from n = 5 experiments and polynomic fit); rather, they decrease slightly, perhaps because of reduced perfusate volume per roller contact with increasing pump rate (rubber tube might not fully snap back into round shape for faster pump rates). **D** Pump switching and pulsations. Test for a possible correlation between relative pulsation slope increases or decreases (bottom axis, pulsation slope change normalized to pump rate before switching) and relative oscillation strength P_0_ (normalized to P_0_ before switching). N = 74 data points from switching between different pump rates in n = 20 NBPs (for an individual example experiment see Fig. 1E). Highly significant positive correlation even though there is a high variance in P_0_ (see also Fig. 5A).

Another prediction is that the engagement of Piezo2 and thus the baroreceptive response amplitude P_0_ should increase with increased pulsation frequency which results in a steeper maximum slope (Fig. 7B), whereas it should be insensitive to static pressure changes because of the fast differential gating, which was already shown by our Windkessel experiments (Fig. 7B, 2B). In these experiments, the pulsation slope is reduced, but not the pump frequency and thus not the static pressure; therefore, the abolishment of the slow LFP oscillation by the Windkessel rules out a substantial role of the static pressure level. The prediction of a positive correlation between pulsation frequency and strength of baroreceptive response was successfully tested via analyses of LFP recordings in which the pump frequency was changed, usually in steps of ± 3 rpm (± 0.4 Hz roller frequency) (n = 20 NBPs, Fig. 7C, D; exemplary experiment shown in Fig. 1E). The overall screening of pulsation frequencies and therewith the tuning of the adequate stimulus was limited on the lower side by the disappearance of PNA and the requirement of pressure levels sufficient for perfusion, and on the upper side by the increasing probability for leaks in both the perfusion system and the vascular system due to the high pressure. In conclusion, Piezo2 is a very plausible substrate for the observed baroreceptive transduction.

Thus, our experiments yield evidence for the ability of physiological vascular pressure pulsations to directly modulate neuronal activity in a subset of neurons in the mammalian brain via mechanosensitive ion channels.

## Discussion

Here we report a direct modulation of the excitability of central neurons in the OB via cardiovascular pressure pulsations that are transduced by mechanosensitive ion channels (summarized in Fig. 8). While increased excitability of central neurons in response to mechanical stimuli has been demonstrated previously *in vitro* in acute brain slices (Nikolaev et al., 2015) and in culture (e.g. Gaub et al., 2020), here we provide evidence for the direct detection of vascular pulsations by neurons within intact brain tissue itself, modulating spiking activity of OB neurons both *in vivo* and in the *in situ* nose-brain preparation (NBP).

**Figure 8:**
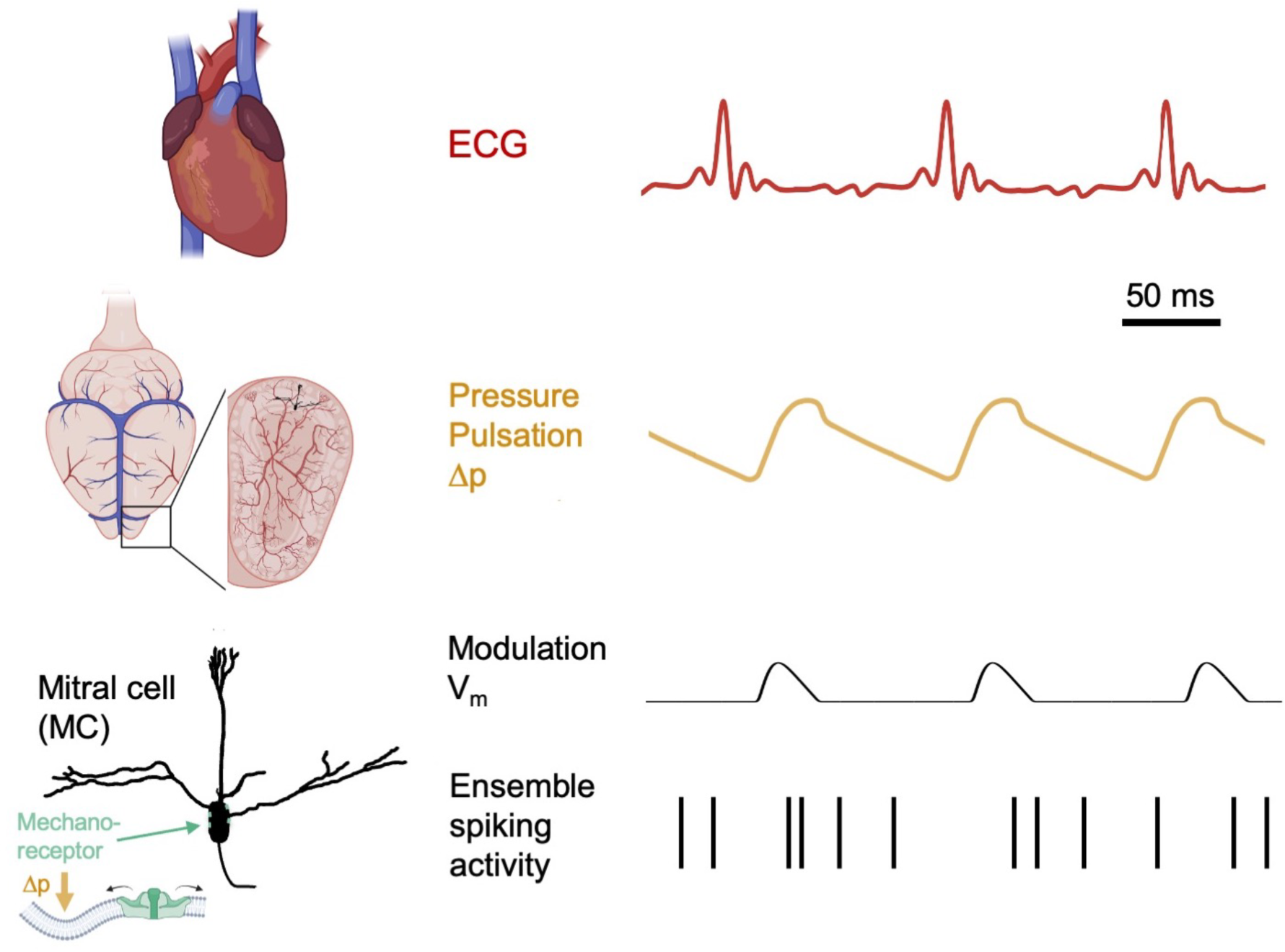
Graphical abstract of main findings Top: Left: rat heart, Right: schematic rat electrocardiogram (ECG). Middle: Left: schematic rat brain and coronal section through olfactory bulb and vessels, exemplary mitral cell (black). Right: intracranial pressure pulsations due to heartbeat. Bottom: Left: Mitral cell from above with mechanoreceptors (green) in membrane (might also be located in dendrites). Right: Modulation of mitral cell membrane potential V_m_ by positive inward currents, below ensuing modulation of spontaneous spikes in mitral cell ensemble. Pressure pulsation trace modified from Reddy et al. 2013. Created in part with www.biorender.com.

The NBP has allowed to reveal this pulse pressure modulation for the following reasons:

1. In this reductionist approach, neural activity is independent of respiratory sensory afferents from the nose because of the absence of nasal airflow, and also not influenced by the massive secondary modulatory inputs to the OB because of the decerebration that disconnects the OB from any other afferent synaptic inputs. These include afferents from the respiratory motor networks in the brainstem (Yang & Feldman, 2018) and from primary olfactory cortices and neuromodulatory centers such as the basal forebrain (reviewed in Brunert & Rothermel, 2021). Accordingly, our *in vivo* recordings from awake mice revealed a lower modulation strength and a much smaller fraction of pulsation-entrained neurons compared to respiratory modulation (110% versus 130% mean modulation index and 10% versus 90 % of recorded units, Fig. 4F, G, Fig. S4A, B). These findings can also explain why the subtle pulsation-mediated modulation of neuronal excitability by heartbeat has not been observed previously.
2. The amplitude and the spectrum of pressure pulsations induced by the peristaltic pump in our system closely resemble those generated by the heartbeat *in vivo*. While in humans arterial blood pressure pulsations Δp_Pulsation_ are known to range on the order of 50 ΔmmHg, intracranial pressure pulsations related to heartbeat are substantially attenuated (reviewed in Wagshul et al., 2011), and a similar attenuation takes place in rats and dogs (Kim et al., 2012); the cardiac component of intracranial pressure pulsations in rat is on the order of ≤ 5 ΔmmHg (Eftekhari et al., 2020) and thus intracranial pressure modulations *in vivo* will be in the range of the magnitudes of the pump-induced pressure pulsations observed in the present study (Fig.2, Fig.7C): When accounting for the drop in pressure across the cannula (see Methods), pulsations at the entry point into the NBP can be expected to occur at ∼ 2 ΔmmHg. Importantly, dampening of these pulsations by a Windkessel device accordingly resulted in the disappearance of the corresponding slow LFP signal (Fig. 2B). Pulsation frequency (as imposed by the tuning process of the NBP, see Methods) was on the order of 2 - 4 Hz; heartbeat frequency in humans is 1 - 2 Hz (60 - 120 bpm) while in rodents it ranges on the order of 5 - 10Hz (300 - 400 bpm for rats, 400 - 700 bpm for mice, e.g. Janssen et al., 2016; Meller et al., 2011; Reddy et al., 2014). Thus, here a common nuisance in perfused preparations can be taken advantage of to explore physiologically relevant phenomena.

However, the pressure pulsations imposed by the pump in our system are at this point limited to a lower frequency band ≤ 4 Hz and also do not provide an exact playback of heartbeat-induced Δp_Pulsation_ in terms of waveform. Although the sinusoidal deflections constitute a decent first-order approximation, the systolic pressure rise during actual heartbeat is much faster than the diastolic decay in both humans and rodents; while there is temporal filtering due to attenuation, the rising phase can still be expected to last less than one third of the entire cycle (see e.g. Fig. 7 in Wagshul et al., 2011, Fig. 1 in Reddy et al., 2014). If the transduction of pressure pulsations into slow OB LFP oscillations is as suggested mainly mediated by Piezo2, which is sensitive mostly to the upward slope of the pulsation (see also Fig. 7 and below), the physiological pulsation transduction should be even more accentuated *in vivo* with otherwise similar parameters (e.g. Δp_Pulsation_ amplitude and frequency).

In the NBP we observed that baroreceptive LFP oscillations originate mostly in the vicinity of the mitral cell layer (see also below). However, the silicon multi-electrode recording approach used for the *in vivo* experiments revealed the presence of heartbeat-entrained neuronal units across all OB layers. This finding could be explained by the fact that the OB contains mostly inhibitory neurons (∼80% of population, Shepherd et al., 2004), that are mostly driven by the principal mitral and tufted cells, the predominant OB excitatory cell type. Thus, we propose that the pressure-entrainment of units in all the other layers is driven by mitral cells, most likely the main pulsation-sensitive neuron population according to our NBP data (Fig. 2B, 5C), via dendrodendritic and axonal contacts to granule cells (Isaacson & Strowbridge, 1998; Pressler & Strowbridge, 2020; Schoppa, 2006), and connections to tufted cells along with possibly other juxtaglomerular neurons via gap junctional coupling (Ma & Lowe, 2010).

### Baroreceptive afferents in the brain versus direct mechanical transduction of pulsations

At this point there is ample recent evidence that heartbeat widely influences neural activity across various brain areas (e.g. Kim et al., 2019; reviewed in Park & Blanke, 2019, see also below). So far, this modulation was proposed to happen mostly via detection of heart pulsations by arterial baroreceptors and cardiac mechanosensory neurons, such as the classical arterial baroreceptor pathway ascending via the Nucleus Tractus Solitarius (NTS, Saper, 2002), and vagal afferents (Karemaker, 2022). From the NTS there are projections to thalamic nuclei which in turn project to viscerosensory cortices including the insula, amygdala and cingulate cortex and are thought to be the main source for heartbeat evoked potentials (HEPs) (Park & Blanke, 2019). Baroreceptor firing can encode both phasic and tonic arterial pressure changes, i.e., both pulsations and changes in mean blood pressure levels. No direct projections from the NTS to the OB are known at this point, however baroreceptive information might reach the OB via limbic afferents or the olfactory tubercle.

In any case, might baroreceptive afferents explain our findings? The observations in the NBP cannot be due to such thalamic relays of the ascending baroreceptor reflex pathway (see Dampney, 1994; Ricardo & Koh, 1978). As discussed above, decerebration physically removes all potential ascending inputs to OB and the fact that the slow LFP oscillations are insensitive to blockade of action potential conduction (Fig. 6B) excludes any contribution of a secondary modulation via afferent baroreceptive inputs from the cardiovascular system. In the *in vivo* experiments, the peak of entrained neural activity at ∼ 20 ms past the ECG peak (Fig. 4D, J) occurs much too early for a visceral ascending pathway involving multiple synaptic relay stations. Indeed, human intracranial HEPs are reported to peak at 200 - 300 ms past the ECG peak (Park & Blanke, 2019). Rather, 20 ms are consistent with the travelling time of the blood pressure wave from heart to brain (∼ 10 ms, see Results) and support a fast mechanotransduction mechanism (see below). In humans, intracranial HEPs due to such a fast component of mechanical modulation of neuron excitability might be masked by the ECG R-peak artefact.

### Mechanotransduction within the mitral cell layer: Evidence and putative mechanism

Several lines of experimental evidence provided in this study suggest that the pulsation transduction is generated via mechanosensitive channels specifically within mitral cell membranes:

1. The slow, pulsation-dependent LFP oscillations observed in the NBP do not require network activity (i.e. are insensitive to the blockade of Na_v_ channels and therewith spiking, Fig. 6B).
2. Mitral cell spontaneous spiking is in part synchronized to the LFP oscillation (Fig. 3), which means that the pulsatile pressure modulation must directly affect the mitral cell membrane potential V_m_ in order to influence their spiking.
3. The origin of the oscillatory signal can be tied mostly to the mitral cell layer (Fig. 5C).

The observation that spiking of mitral cells is synchronized to the upward phase of the oscillation with respect to the mitral cell membrane potential (Fig. 3B) indicates that the transduction must be excitatory, i.e., via influx of cations.

Throughout the body, a huge diversity of mechanosensitive ion channels mediate a wealth of exteroceptive (tactile, auditory and vestibular perception, nociception) and interoceptive sensory modalities, linked to the sensing of pressure changes and/or membrane stretch in e.g. the arterial vasculature, lungs or gastrointestinal system (reviewed in Jin et al., 2020). Mechanosensitive K^+^-specific channels such as K2P (K^+^ two-pore) channel types can be ruled out because of the excitatory nature of the pulsation transduction mechanism in mitral cells, and the blockade of LFP oscillations by the tarantula venom GsMTx4 (Suchyna et al., 2000, 1.25 µM D-GsMTx4) provides proof for cationic mechanosensitive channels such as Piezo channels, TRPC1 and TRPC6 channels and TACAN (Tmem120a) (Beaulieu-Laroche et al., 2020; Suchyna, 2017).

Due to their extremely fast gating (activation < 1 ms), the Piezo channel family (Piezo1 and Piezo2) has been identified as a key element of fast mechanical signal transduction in arterial baroreceptors (Zeng et al., 2018), lung stretch receptors (Nonomura et al., 2017), outer hair cells (Li et al., 2021) and in a variety of skin (tactile vibration) receptors as well as proprioceptors (reviewed in (Anderson et al., 2017). So far, Piezo2 is the channel that is predominantly expressed in sensory and proprioceptive neurons whereas Piezo1 is found in other cell types such as endothelial cells and muscle cells, but there is also overlapping expression (reviewed in Fang et al., 2021; Murthy et al., 2017).

The most compelling evidence for Piezo2-driven mechanotransduction in the mitral cell layer is provided by recent reports that showed the specific expression of Piezo2 in a subset of mitral cells within the OB (but not or barely within other OB cell types) at the RNA and protein level (Wang & Hamill, 2021; Zeppilli et al., 2021). Moreover, the transformation of pressure pulsation into LFP waveform is best explained by Piezo2 properties, rather than Piezo1, due to the rapid inactivation of Piezo2 and its preferred activation by pressure increase (Fig. 7A, e.g. Coste et al., 2010; Lewis et al., 2017). Fast inactivation would allow the response to follow the pressure pulsations with almost no filtering, as evident from the highly symmetric LFP waveform reconstructed from the oscillation harmonics (Fig. 2E, 7A). Some multimodal TRP channels may also be gated by mechanical stimuli (Liu & Montell, 2015); mechanosensitivity has been ascribed to TRPC1 and TRPC6 (e.g. Alessandri-Haber et al., 2009). However, blockade of TRPs had no effect on the baroceptive response (Fig. 6D). Moreover, according to Zeppilli et al., (2001, online database) TRPC6 is not expressed in mitral cells (but in external tufted cells), there is weak expression of TRPC1 in a subset and finally, Piezo1 and TRPV4 are not expressed. In conclusion, Piezo2 emerges as a highly likely candidate for the molecular correlate of rapid pulsation transduction in mitral cells observed here. Since we cannot exclude the involvement of other mechanosensitive channels, the exact substrate of mitral cell pulsation transduction remains to be elucidated via studies at the single cell level.

### Possible link to respiratory rhythm, coupling between respiration and heartbeat

So far, respiration is well known to generate a distinct neural rhythm that is detectable across almost every brain structure. In the OB the respiratory rhythmicity is functionally imposed by the temporal variation in odor sampling during nasal respiration. Beyond this straight-forward coupling between the rhythmic presence of the odor stimulus and olfactory sensory neuron activity, and the ensuing modulation of second order neuron activity in the bulb (e.g. Buonviso et al., 2006; Rojas-Líbano et al., 2014, our Fig. S4), nasal mechanoreceptors and trigeminal afferents also contribute to this patterning even in the absence of odorants (Aucoin et al., 2023; Grosmaitre et al., 2007). Respiratory rhythmicity is transmitted to upstream higher brain areas such as the hippocampus, prefrontal cortex and primary visual cortex (reviewed in Goheen et al., 2023; Tort et al., 2018); there respiration patterning most likely arises from efference copies from the respiratory centers within the brain stem (Karalis & Sirota, 2022).

The neural heartbeat detection mechanism in mitral cells described here might also contribute to an intrinsic resonance of the bulbar and other central networks to respiration, since breathing-related intracranial pressure changes are in a similar amplitude range as heart-beat mediated pulsations have been reported (as proposed by Wang & Hamill, 2021, their Fig. 7). Respiration in turn is interlinked with heartbeat in various ways, most prominently via the phenomenon of human heart rate variability that is directly imposed by the breathing rhythm and mediated by the vagus nerve (reviewed in Karemaker, 2022). While heartbeat-dependent modulation of neural activity in the OB did not require nasal airflow in our *in vivo* experiments (Fig. S4), more subtle interactions cannot be excluded. In rats, high-frequency sniffing in otherwise resting animals increases heart rate to values similar to those in motion periods (Kuga et al., 2019).

Finally, beyond its expression within olfactory bulb mitral cells, Piezo2 has been observed to be expressed in subsets of principal neurons across various brain areas, the hippocampus, neocortex and cerebellum (Wang & Hamill, 2021), that might all serve as interoceptive “sentinel neurons” for both heartbeat and respiration induced pressure pulsations.

### Possible physiological function(s): Interoceptive modulation

Interoception transmits various bodily states to the brain, whereby autonomic tone can modulate exteroception (i.e., sensory perception of the outer world) and cognition. Aside from the respiratory and cardiac modulations discussed above, another more recently uncovered example is the vegetative nervous system of the gastro-intestinal tract which generates peristaltic gut movements which in turn can modulate cortical activity in a wide-spread fashion (Rebollo & Tallon-Baudry, 2022).

While the physiological relevance of the so far little known autonomous interface between body and mind is poorly understood at this point, various hypotheses are under debate (see e.g. Tallon-Baudry, 2023).

1. Interoception of rhythmic processes might confer some type of clocking and/or global pacemaking and thus allow to coordinate neural activity across brain areas on intermediate timescales in the delta and theta range (as compared to gamma rhythms that are also implied in binding activity).
2. Interoception might be related to resting states. For example, respiratory coupling of cortical oscillations is unmasked during sleep (Karalis & Sirota, 2022). Here an experimentally reductionist approach has allowed to reveal pressure-pulsation mediated coupling. While these observations seem to imply that interoception is more prominent at rest, it might be simply easier to detect because of reduced overlap with other activity such locomotion which is also correlated with massive multisensory input.
3. As to the mechanism for heartbeat transmission reported here, fast excitatory feedback for an increased heart rate (vs. slower feedback due to the classical baroreflex) might facilitate adequate responses due to arousal occurring during e.g., the response to predators or social interactions. Arousal and attention-demanding tasks are known to activate cholinergic projections (reviewed in Sarter et al., 2005) which in turn act both in a vasodilative manner to increase blood flow (Sato & Sato, 1992) and to directly modulate neuronal activity, which was previously shown by us to be correlated with OB-based social discrimination (Suyama et al., 2021). Such coincident effects – a direct increase in excitability via fast-acting mechanosensors within the OB <u>and</u> top-down release of neuromodulators – might further summate in order to increase e.g. sensory acuity during a relevant context via modulation of brain regions that are not targeted by the classical baroreceptive afferents.
4. (4) Emotional processing is known to involve an autonomous component which might involve respiration, gut movements as well as the cardiovascular system, leading back to the somatic marker hypothesis by which emotional states and conscious awareness are a consequence of interoception (Candia-Rivera, 2022; Damasio, 1999). Recently, optogenetically increased heart rates in mice were shown to recruit anxiogenic brain circuits (Hsueh et al., 2023), further enhancing the plausibility of an interoceptive modulation of central neural circuits.
5. (5) In humans, there is strong evidence with regard to the influence of heartbeat cycle, as detected via heartbeat evoked potentials (HEPs) in the brain (reviewed in Coll et al., 2021), on both perception and cognition. For example, heartbeat modulates somatosensory acuity and self-face recognition (Al et al., 2020, 2021; Sel et al., 2017), and even racial stereotype expression has been shown to synchronize with heart cycle (Azevedo et al., 2017).

The physiological relevance of heartbeat mechanosensing in the OB is far from clear at this point, nevertheless the results of the present study have unraveled a hitherto unknown, brain-intrinsic mechanism for direct detection of cardiovascular activity (compared to the known baroreceptor and cardiac afferents). Considering the recent evidence for a distributed expression of mechanosensitive neurons in the brain it is tempting to speculate that mechanosensitive central neurons could form a network of “sentinel neurons” that can contribute to interoceptive modulation of cognition, mood and autonomic status itself.

## Acknowledgments

We wish to thank Anne Pietryga-Krieger for technical assistance, Günes Birdal for stainings, Alice Stephan for help with coding, Esteban Pino for help with pressure recordings, and Dres Mohammad Farman Tariq, Owen Hamill, and Frank Schweda for discussions and/or input on the draft. This work was supported by DFG-EG135/7-1 and FOR5424 (VE).

## Author contributions

LJS performed all the nose brain preparation experiments with initial help from MD and the basic analyses of the OB-LFP recordings and contributed to the conception of experiments and to writing of the Methods section. SHB performed all the in vivo experiments and their analyses (Fig. 4, S4) and wrote the respective parts of the manuscript. ILHO contributed to the conception of *in vivo* experiments. VE guided the study, contributed to coding of the NBP analyses, conceived and performed meta-analyses of NBP data and wrote the manuscript except for the abovementioned parts. All authors participated in the final phase of manuscript editing.

## Glossary

ECG: electrocardiogram
FFT: Fast Fourier Transform
HEP: heartbeat evoked potential
LFP: local field potential
MCL: mitral cell layer
NBP: nose-brain preparation
NTS: Nucleus tractus solitarius
OB: olfactory bulb
PNA: phrenic nerve activity

## Notes

### Competing Interest Statement

The authors have declared no competing interest.

