## SupplementaryMaterial for "Blood pressure pulsations modulate olfactory bulb neuronal activity via mechanosensitive ion channels"

### Supplementary Material

Numbering indicates association with respective Figure in Results part, except for Supplementary Table 1 (which belongs to the Methods part)

#### Supplementary Table S1 Animals

| Strain | Wild type Wistar | VGAT Venus | VP-EGFP | TH-GFP |
|---|---|---|---|---|
| N animals total | 102 | 5 | 20 | 15 |
| N WT/N transgenic same period, with detectable $P_0$ | | 15/5 | 28/14 | 29/14 |
| Mann-Whitney test result normalized $P_0$ amplitude | | 0.294 | 0.298 | 0.66 |

**Supplementary Table S1: Experimental animals and comparison between experiments from WT and transgenic strains.**

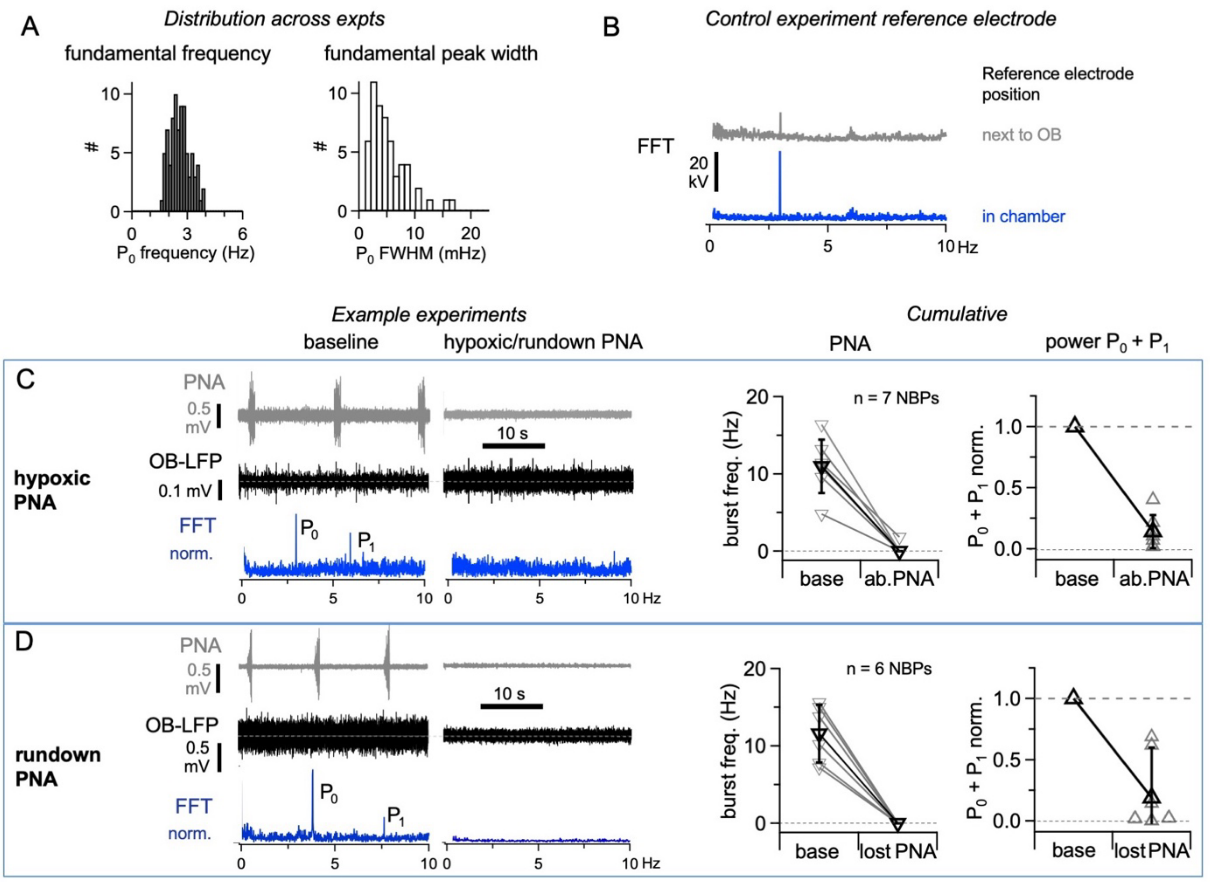


**Supplementary Figure S1: LFP oscillations are independent of the reference electrode positioning and require viable neuronal tissue.**

**A** Left panel: Distribution of $P_0$ frequency in our initial set of experiments (mean 2.6 ± 0.5 Hz, range 1.5 - 4 Hz, n = 83 NBPs) before the relation between LFP oscillations and pump rate was revealed. The apparently biological distribution is due to the fact that each preparation undergoes an individual tuning of the optimal flow and thus pump rate. Right panel: Distribution of the full width half maximum (FWHM) of $P_0$ in the same set of experiments. The FWHM determination was sometimes limited by the FFT scaling $\Delta x_{FFT}$ which for the most commonly used LFP data interval of 360 s sampled at 1 kHz amounted to $\Delta x_{FFT}$ = 2*500 Hz/360000 = 2.8 mHz. Thus the tuning is likely narrower than reported here.

**B** Control for LFP oscillations with differing position of reference electrode. FFTs of recordings next to OB (top) and outside within recording chamber.

**C** Set of experiments in which hypoxia was induced by lowering the pump rate to 3 rpm for 10 minutes. Left panels: Example experiment recordings and FFT of OB-LFP before (baseline, left) and after hypoxia induction and reinstatement of the original pump rate (right; FFT spectra normalized because recordings were of different duration). Right panels: Cumulative data of all n = 7 experiments. PNA burst frequency, and $P_0 + P_1$ amplitudes normalized to the baseline value. Baseline mean PNA frequency 11.0 ± 3.5 bursts/min, hypoxic condition 0.0 ± 0.0 bursts/min. Mean power amplitude ($P_0 + P_1$) in hypoxic condition 0.14 ± 0.13 of baseline, $P < 0.01$, Wilcoxon test.

**D** Set of 6 experiments in which there was eventual rundown of PNA. Panels as in C. Baseline mean PNA frequency = 11.6 ± 3.7 bursts/min, rundown condition 0.3 ± 0.7 bursts/min. Mean $P_0 + P_1$ amplitude in rundown condition = 0.19 ± 0.26 of baseline.

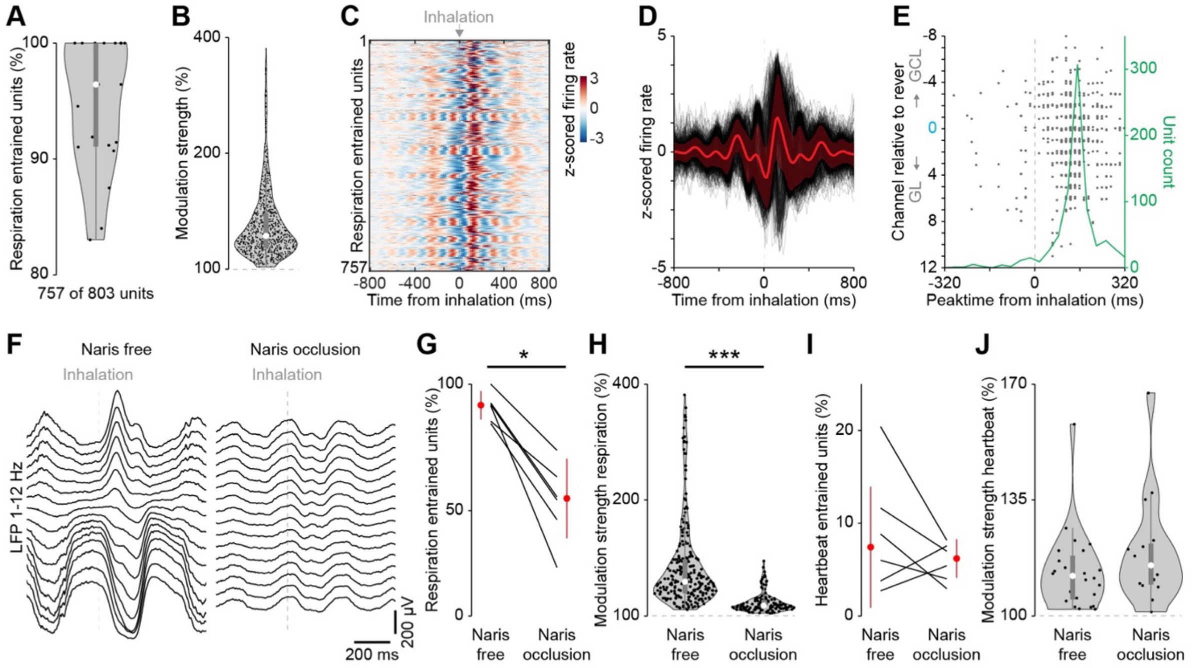


**Supplementary Figure 4: Respiratory modulation of bulbar neurons *in vivo* and effects of naris occlusion on respiratory entrainment and heartbeat entrainment**

Same set of experiments as in Fig. 4.

**A** Percent of respiration entrained units for 19 recording sessions from 11 mice.

**B** Modulation strength of respiration entrained units.

**C** Color-coded z-scored firing rate of significantly entrained units aligned to respiration.

**D** Z-scored firing rate of significantly entrained units aligned to respiration. Red is the average of all significant units.

**E** Distribution of peak firing rate time points of respiration entrained units versus channel number relative to the reversal channel (blue 0) and the corresponding histogram (green).

**F** Example LFP (1-12 Hz) aligned and averaged for 1000 inhalations for all channels without and with naris occlusion.

**G** Percent of respiration entrained units for 6 recording sessions from 6 mice without and with ipsilateral naris occlusion.

**H** Modulation strength of respiration entrained units without and with ipsilateral naris occlusion.

**I** Percent of heartbeat entrained units for 6 recording sessions from 6 mice without and with naris occlusion.

**J** Modulation strength of heartbeat entrained units without and with naris occlusion.

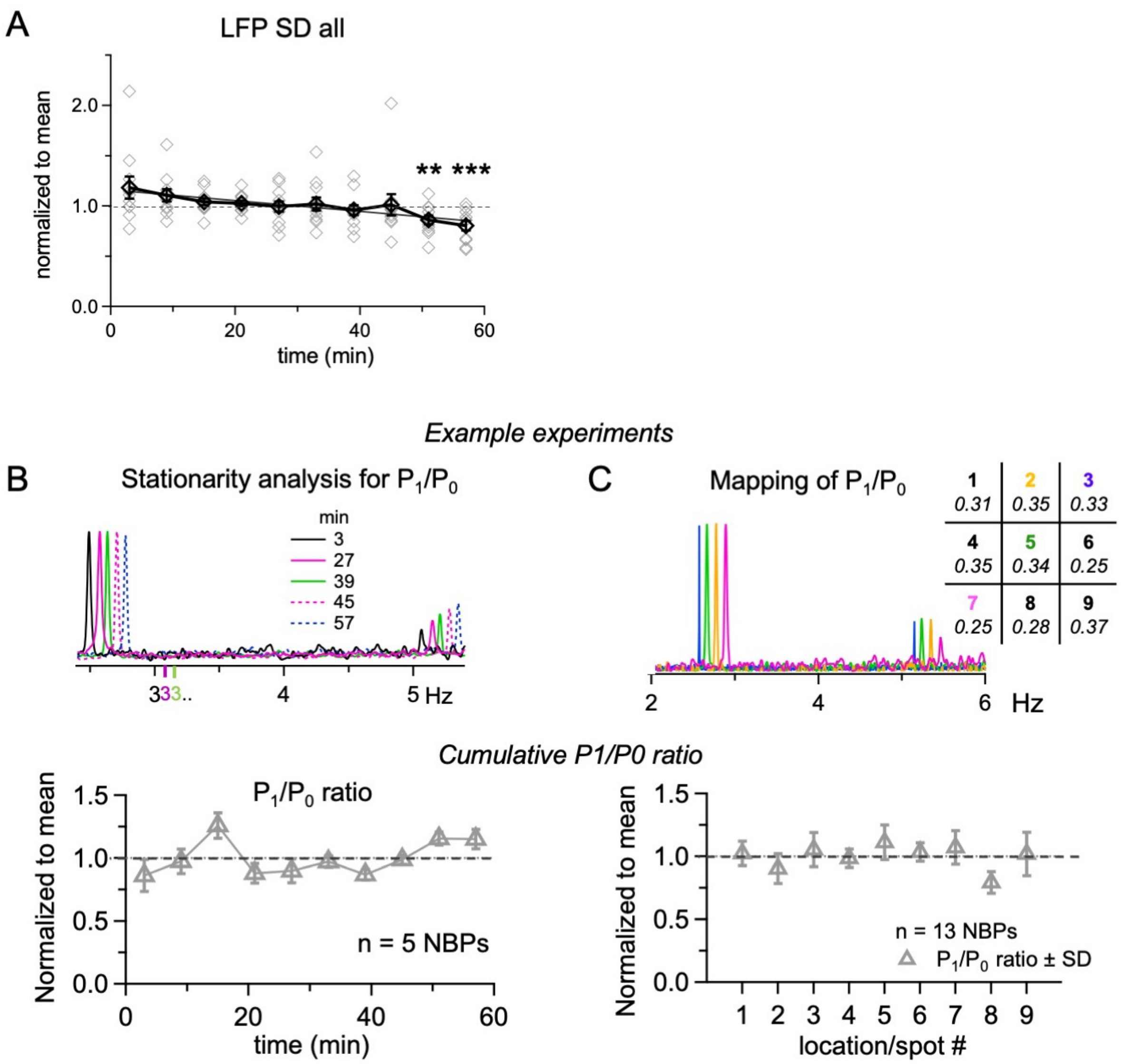


**Supplementary Figure S5: Stability of the LFP activity and harmonics over time.**

**A** LFP standard deviation analysed over time as in Fig 5A. There was a weak, but significant rundown over time (linear fit slope = -0.5%/min). This decline is commonly observed in long-term recordings in the perfused brainstem preparation. It is most likely linked to progressive widening of the interstitial space in response to ongoing retrograde perfusion of the brain tissue (M. Dutschmann, personal communication).

**B** Fraction of first harmonic $P_1$ relative to $P_0$ analysed over time as in Fig. 5A. Top: FFT set of example experiment; frequencies across time points were identical but shifted in order to allow for better visual comparison. Bottom: Cumulative data; individual data points not shown. Deviations from the mean were not significant.

**C** Fraction of first harmonic $P_1$ relative to $P_0$ analysed over space as in Fig. 5B. Top: FFT set of example experiment; the frequencies across locations were identical but shifted in order to allow for better visual comparison. Bottom: Cumulative data; individual data points not shown. Deviations from the mean were not significant.

### Supplementary Table S6 Pharmacology

| Condition / parameter | Control (Fig. 5A) | ACSF injection | Lidocaine injection | D-GsMTx4 injection | SKF injection |
|---|---|---|---|---|---|
| N experiments | 10 | 6 | 8 | 13 | 12 |
| $P_0 + P_1$ power | 94 ± 43 | 91 ± 46 | 130 ± 88 | 13 ± 11*** | 106 ± 36 |
| Spike count 2π | 97 ± 44 | 77 ± 44 | 25 ± 33** | 94 ± 24 | 93 ± 17 |
| LFP SD | 83 ± 20 | 86 ± 37 | 63 ± 33** | 125 ± 59 | 100 ± 16 |

**Supplementary Table S6: Controls and pharmacology**

Mean values and SDs of all cumulative parameter plots shown in Fig. 6, and for time points 2 and 7 from the stationarity test shown in Fig. 5A/S5A (Control).
